# The inter-continental population dynamics of *Neisseria gonorrhoeae*

**DOI:** 10.1101/2023.08.02.551601

**Authors:** Magnus N Osnes, Ola B Brynildsrud, Kristian Alfsnes, Lucy van Dorp, Samantha A McKeand, Jonathan Ross, Katy Town, Jolinda de Korne-Elenbaas, Alje van Dam, Sylvia Bruisten, Birgitte F de Blasio, Dominique A Caugant, Yonatan H. Grad, Deborah A Williamson, Francois Balloux, Xavier Didelot, Vegard Eldholm

## Abstract

*Neisseria gonorrhoeae* is a globally distributed sexually transmitted bacterial pathogen. Recent studies have revealed that its evolution has been shaped by antibiotic use, while molecular surveillance efforts have demonstrated large changes in lineage composition over relatively short time-spans. However, the global population dynamics of *N. gonorrhoeae* remain unsatisfyingly characterized.

To reconstruct recent large-scale population dynamics, we generated a dated phylogeny from 9,732 *N. gonorrhoeae* genomes and found the effective population size of the species to have expanded gradually over the past two centuries. While the effective population size of clades with reduced susceptibility to extended-spectrum cephalosporins started declining around 2010, a major clade containing a mosaic *mtr* operon associated with cephalosporin susceptibility and decreased azithromycin did not display any reduction in population size.

Using ancestral trait reconstruction analyses, we delineated transmission lineages, defined as groups of sequences in which all the samples can be traced back to the same import event to a given location. Import, export and local transmission dynamics across two densely sampled locations (Norway and Victoria, Australia) were investigated in detail. Norway exhibited substantially higher rates of strain import and export compared to Victoria, where incidence was to a larger extent fuelled by locally transmitted lineages. Taken together, our work highlights the power of large-scale phylogenomic analyses to uncover the complex dynamics of lineage transmission in *N. gonorrhoeae*.

## Introduction

*Neisseria gonorrhoeae* is a globally distributed obligate human pathogen composed of lineages undergoing rapid fluctuations in prevalence. A prominent example is multilocus sequence type (ST) 1901 which was shown to have undergone dramatic fluctuations in effective population size (1) concomitantly with changes in prevalence and sexual network association (2). Analyses of a global collection of 419 gonococcal genomes demonstrated that antibiotic use likely played a central role in shaping the recent evolution of *N. gonorrhoeae* (3). The study also suggested an African or European origin of *N. gonorrhoeae* and that the species primarily expanded in Asia.

As no gonococcal vaccine is currently available, public health management of gonorrhea largely relies on empiric diagnosis and antibiotic therapy. It is thus concerning that gonococci have evolved resistance to all antibiotics recommended to treat gonorrhea. Around 2012, cefixime monotherapy was replaced by dual therapy consisting of ceftriaxone and the macrolide azithromycin in most of Europe (4) and the USA (5). Both cefixime and ceftriaxone are third generation cephalosporins, but mutations reducing susceptibility generally have limited effect on the minimum inhibitory concentration (MIC) of ceftriaxone compared to cefixime. In the United Kingdom (UK), treatment guidelines in 2018 recommended ceftriaxone monotherapy, at a higher dose, reflecting increasing numbers of azithromycin non-susceptible infections (6), with similar updates subsequently implemented in the USA (7), Norway, and elsewhere. The main determinant of cephalosporin resistance is *penA*, and the majority of alleles responsible for reduced susceptibility are mosaic alleles resulting from recombination with other members of *Neisseria* spp. (8, 9).

Mosaic alleles generated by interspecies recombination are also key effectors of reduced susceptibility to azithromycin. High-level azithromycin resistance is typically associated with mutations in *rrl*, encoding the 23S ribosomal rRNA (10). In addition, mutations in *rplD* (*10*), *rplV* (11), the *mtr* operon, the *mtr* regulator *mtrR,* as well as its promoter (12), can synergistically modulate susceptibility and, in combination, lead to azithromycin non-susceptibility. In isolation however, these mutations rarely result in a clinically significant reduction in azithromycin susceptibility. Recently, mosaic alleles spanning the *mtr* operon were shown to cause azithromycin resistance, an effect ascribed to epistatic interactions between *mtrR* promoter mutations and the *mtrCDE* efflux pump (13). The circulation of significant numbers of clinical isolates harboring mosaic *mtr* and exhibiting low-level azithromycin resistance was first detected in the state of Victoria, Australia (14). A subsequent study of a large global collection of gonococcal genomes suggested that the clade harboring mosaic *mtr* had expanded since at least 2011 (15).

Globally, the evolution of *N. gonorrhoeae* has been shaped by antibiotic use (1, 3), while molecular surveillance efforts have demonstrated large-scale changes in lineage composition over relatively short time-spans (2). There have been studies investigating the international dissemination of *N. gonorrhoeae* (*2, 16, 17*), but migration dynamics have not been characterized in a quantitative manner. Our current understanding of disease imports, and their importance for country-level epidemiology, is ultimately based on travel histories as stated by individual patients, if known at all. A quantitative understanding of export dynamics is lacking, both due to a lack of available tools and because it falls outside the responsibility of national public health agencies monitoring infectious diseases within country borders.

Here, we used a large collection of genome sequences from clinical isolates to study *N. gonorrhoeae* population dynamics and patterns of dissemination. We generated a dated phylogeny after using a novel and computationally efficient approach to mask the majority of recombined regions. By integrating genomic and epidemiological data, we 1) reconstructed the expansion and contraction of clades harboring genetic determinants of reduced susceptibility to key antibiotics; 2) reconstructed large-scale lineage dynamics across locations ranging in scale from continent to state; and 3) estimated import and export dynamics in two well-sampled locations. Our approach, combining computationally efficient approaches to account for recombination, generate temporal phylogenies, estimate ancestral states, and summarize outputs, represents a useful framework for future large-scale genomic epidemiology studies of microbial pathogens (18).

## Results

### Global population dynamics

To characterize quantitative aspects of gonococcal population dynamics and transmission within and between world regions, we assembled a dataset of 9,732 *N. gonorrhoeae* genomes. To perform detailed analyses of international migration and transmission dynamics, we endeavored to obtain high sampling rates in recent years for countries where this was possible. Reflecting available samples and metadata, the country- and state-level analyses focus on Victoria, Australia, with 2,203 genomes collected in 2017,(14) and Norway, with 1,724 genomes collected in 2016-2018. In both Norway and Victoria, all culture positive cases diagnosed in the period were sequenced, and the collections are thus characterized by minimal bias. Through the sampling periods, the sequenced isolates represented 28.1% and 41.5% of notified cases in Victoria State (14) and Norway, respectively. The geographic distribution of sampled genomes and their phylogenetic relationships are illustrated in Fig. 1. An overview of sampling over time as a function of geographic location is available as supplementary material (Fig. S1).

**Figure 1.**
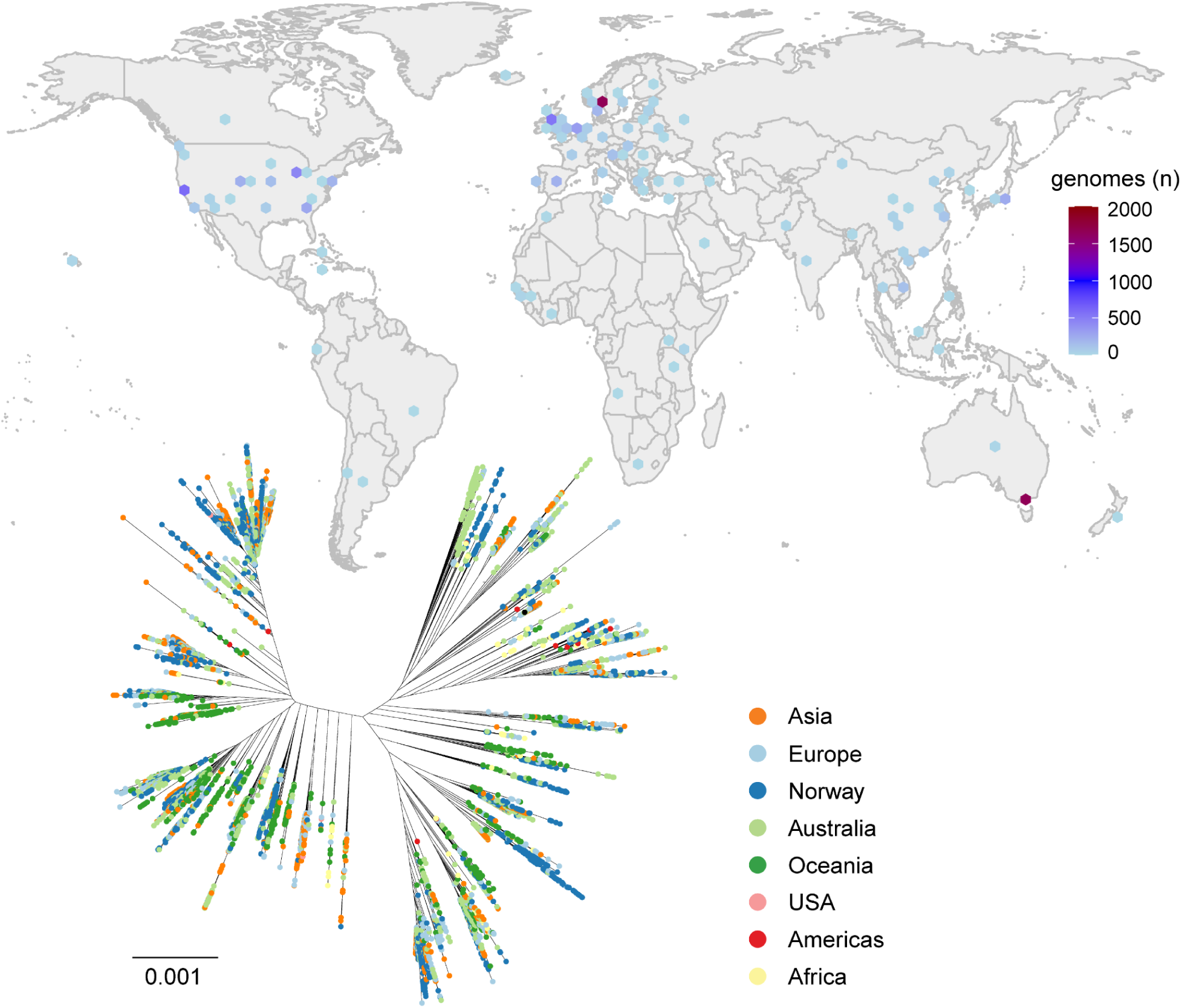
Overview of the sample collection. Top panel: The color of the dots indicate the number of sequences at each location. Bottom panel: Unrooted phylogeny built from a multisequence alignment with partially masked recombinant regions (see main text for details). The tip colors indicate geographic locations. The scale bar represents the number of substitutions per site.

As standard methods to identify recombination events such as Gubbins (19) and ClonalFrameML (20) are unable to handle datasets of this size, we used a novel approach to mask the majority of recombination in the genome dataset (see methods). Next, a dated phylogeny was generated from an alignment of the 9,732 genomes. The global *N. gonorrhoeae* effective population size through time, reconstructed from the dated phylogeny using skygrowth (21), was found to have expanded gradually over the last two centuries. Since there is no indication of a reduction in actual gonorrhea case numbers in the period 2010-2018 (22, 23), an inferred dip in the effective population size in the last decade was unexpected. We reasoned that the inferred dip might be an artifact resulting from non-random sampling. Particularly the sequences from the USA (2016–2018), mainly sampled as part of the GISP and eGISP (24), are enriched for non-susceptible isolates by design. Such sampling bias could affect estimates of effective population size, through undersampling of the true genetic diversity (25). To investigate whether the recent reduction in effective population size may be caused by sampling biases, we generated a phylogeny from subsets of the sequence collection characterized by unbiased sampling schemes. This subsample included 4,184 genomes from Norway, Australia (14), Denmark (26) and the ‘Gentamicin for the treatment of gonorrhea’ trial in the UK (27). Historical genomes from Denmark were primarily retained to ensure that temporal calibration points were available for the generation of a dated phylogeny, whereas the other studies were characterized by inclusive sampling. When the effective population size through time was reconstructed from this dataset (see methods), the dip since ∼ 2010 was far less pronounced (Fig. S2), indicating that the drastic reduction in effective population size inferred from the full sample collection was partially due to sampling bias.

In addition to overall population dynamic patterns, we were particularly interested in clades harboring alleles associated with reduced susceptibility (RS) to ceftriaxone, cefixime, and azithromycin. For azithromycin we focused on mosaic *mtr* alleles, as strains harboring these are associated with elevated azithromycin MICs and have successfully generated large transmission clusters (14). In contrast, mutations in *rrl* were exclusively found as singletons or in small clusters in the phylogeny (Fig. S3), in line with findings from a population-level analysis from Norway (28), and were thus deemed less relevant in this context.

To identify mosaic *mtr* alleles, sequences corresponding to the *mtrCDE* operon were extracted from the full alignment and analyzed using fastGEAR (29). The analysis revealed the presence of five major *mtr* lineages, of which four were clearly mosaic based on visual inspection of the alignments. All the mosaic alleles were mapped to a large monophyletic clade dominated by ST-9363 (Fig. 2). Isolates harboring mosaic *mtr* alleles exhibited elevated azithromycin MICs, with individual *mtr* lineages exhibiting median MICs between 0.5 and 2 μg/ml (Table S1, Fig. S4), compared to 0.125 for isolates carrying non-mosaic *mtr* alleles.

**Figure 2.**
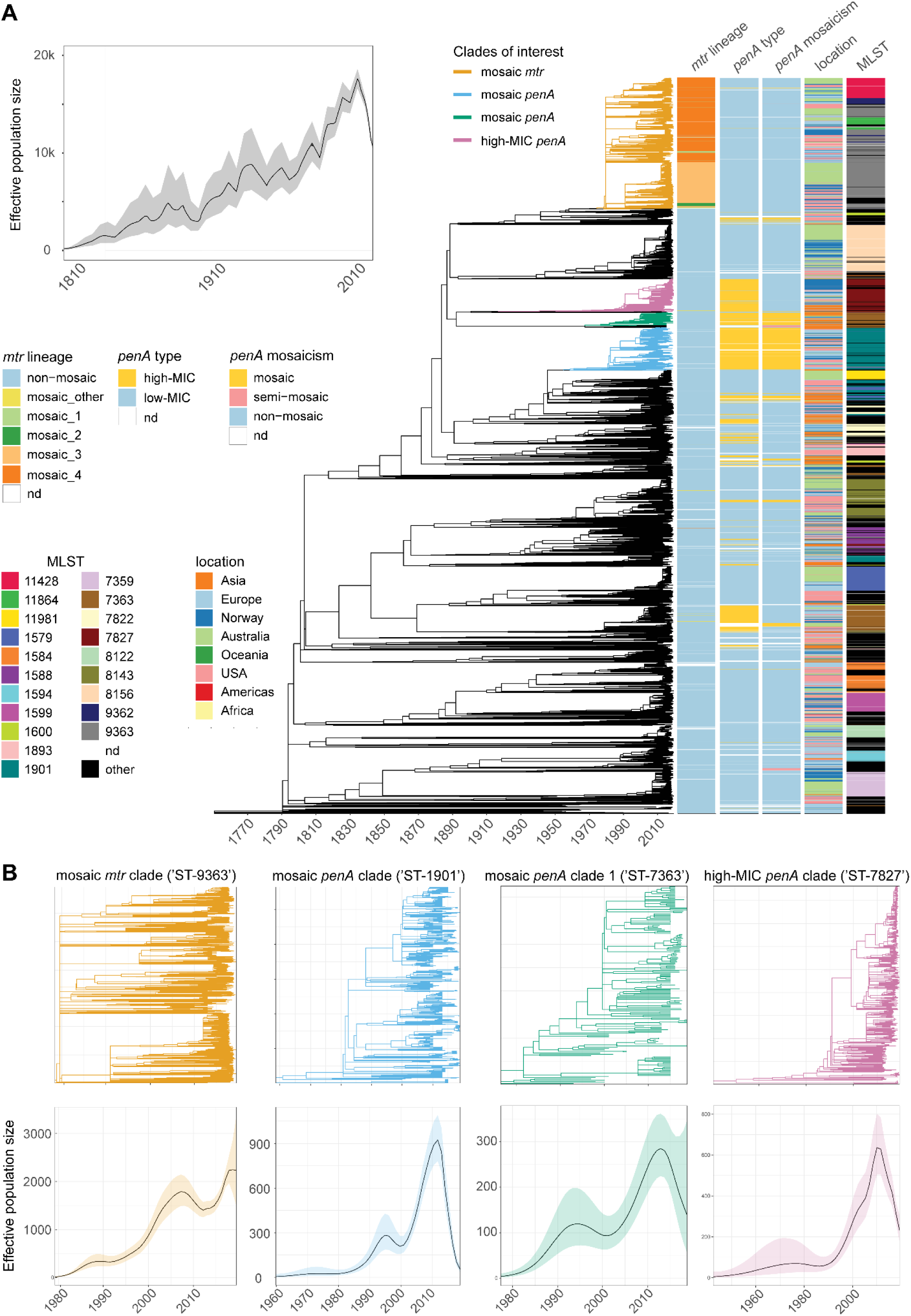
Dated phylogenetic analyses. A) Reconstructed effective population size over time and full tree including annotated metadata. Annotations from left to right: mtr fastGEAR defined allele, ceftriaxone MIC category (blue low, red high), penA mosaic (blue no, orange yes), geographic region. Selected clades in the tree associated with mosaic mtr or penA operons, or associated with high cephalosporin MICs (alleles associated with cefixime and/or ceftriaxone MIC ≥ 0.064) are highlighted with different colors. FastGear identified five mtr lineages, of which 1-4 were mosaic whereas lineage 5 was non-mosaic. Mosaic alleles exhibiting significant sequence variation relative to the major lineages are annotated simply as ‘mosaic’. B) Effective population size for four clades of interest. The coloring corresponds to the clades in the full tree.

Next, we identified clades harboring *penA* alleles associated with elevated cephalosporin MIC. The largest clades carrying *penA* RS alleles were dominated by ST-1901, ST-7363, and ST-7827. Within each clade, the dominant *penA* alleles were 34.001, 10.001 and 13.001, respectively (Fig. 2A). The emergence of these sequence types and *penA* alleles have all been described individually in recent years (1, 30, 31).

We then reconstructed the effective population size through time for each of the four large clades harboring *mtr* and *penA* alleles, associated with reduced susceptibility to azithromycin and cephalosporin (Fig. 2B). The clades harboring *penA* RS alleles were observed to have undergone recent declines in effective population size. This finding is in line with earlier observations for one of the three clades (dominated by ST-1901) which exhibited a striking drop in effective population size from around 2010 (1). However, the large clade associated with various versions of a mosaic *mtr* operon, corresponding to the ST-9363 core-genogroup (32), did not display a similar decline in effective population size.

### Regional epidemiology, import and export dynamics

To investigate import, export and transmission dynamics in the four well-sampled locations of interest (the USA, Europe, Norway or Victoria, Australia), ancestral geographic states were inferred (33) separately for each location, using binary categories. That is, we carried out four independent analyses in which a singular location of interest was treated as one geographic state, and all other locations as the alternative state (‘rest of the world’). Sequences were assigned to the same transmission lineage if they were mapped to the same geographical locations without any estimated geographical transitions on the path that connects these sequences in the phylogeny. Summary statistics and visualizations were generated using LineageHomology (34). With this approach, each genome sampled in one of the locations of interest, was assigned to a transmission lineage or identified as a singleton introduction (see methods for details). An interactive project website for the exploration of phylogenetic inferences, transmission lineages, and metadata, is available on GitHub (https://magnusnosnes.github.io/10000_Ngon_genomes/).

For all locations, we estimated that the largest transmission lineages accounted for a high fraction of the local cases. In Norway, Australia, Europe, and the USA, the ten largest transmission lineages accounted for 44.4% and 59.8%, 71.6%, 54.5% of the cases, respectively, whereas singletons (introductions not leading to detected local transmission) accounted for 15.7%, 7.5%, 4.8%, and 6.5% of the cases, respectively. We estimated that the sizes of the transmission lineages were compatible with being drawn from a power-law distribution, Kolmogorov-Smirnov test (p>0.05) for all locations. The estimated power law coefficients alpha were 2.03, 1.86, 1.88, 1.79 for Norway, Victoria, Europe and the USA, respectively.

Imports constituted a major source of new gonorrhea cases in all locations (Fig. 3). In the densely sampled locations, Norway and Victoria, 27.0% and 12.7% of cases were inferred to be new imports, respectively. The high importance of repeated imports on the Norwegian gonorrhea epidemiology was also evidenced by a large number of small and recently imported transmission lineages (Fig. 3B). Europe and the USA were characterized by larger and older transmission lineages compared to Norway and Victoria, which is to be expected as a result of their much larger sizes, both in terms of population and geography, as a relatively larger number of transmissions will be inferred within borders when the region of interest is larger.

**Figure 3.**
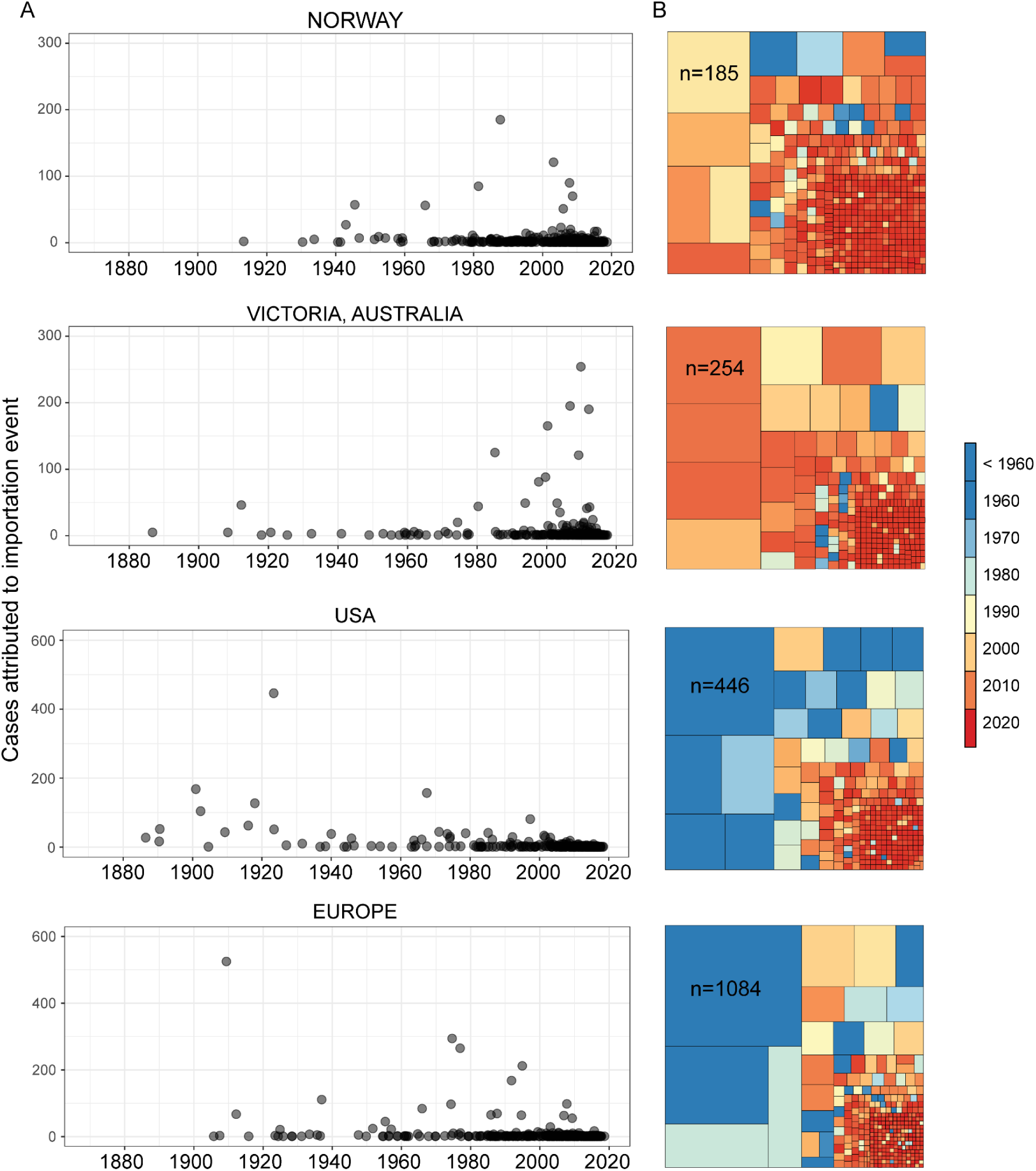
Gonorrhea import dynamics across major sampling locations. A) Transmission lineage size by the inferred times of importation. The largest transmission lineage in Europe (n=1084), was inferred to date back to 1750 and is not shown to maximize resolution. B) Treemaps illustrate the size and age of individual transmission lineages. The size of the squares reflects the relative sizes of the transmission lineages. The color indicates the inferred dates of importation.

In addition to imports, we were particularly interested in export dynamics, as this is not typically considered by public health authorities, and is much harder to assess using classical epidemiological approaches. By dividing the estimated number of exports by the total number of cases in each location, we estimated that each case in Norway and Victoria produced on average 0.22 and 0.07 exports, respectively, indicating substantial variation between locations.

It is worth noting that whereas Victoria and Norway were very densely sampled (all culture positive cases in 2017 and 2016-2018, respectively), this was not the case for the USA and Europe. Sampling density and bias will be a key determinant of quantitative import and export estimates (Fig S6).

To evaluate whether the sample collections were suitable for accurate relative quantification of imports, exports and local transmission in each location, we separately upsampled tips in the dated phylogeny from the focal location and the ‘Rest of the world’ (ROW). We continued the upsampling until all available samples were included (see methods). For each subset, the phylogeographic mapping, transmission lineage assignment and import, export, and local transmission quantification procedure was repeated independently. We reasoned that there was sufficient phylogeographic signal in the datasets if the estimated import (or export) fractions converged at a similar value when: 1) increasing the number of tips included from the focal location, while keeping all tips from ROW, and 2) increasing the number of tips from ROW, whilst keeping all tips from the focal location. If these values did not converge towards similar values, this indicated that one of the categories (focal location or ROW) was insufficiently sampled.

For Norway, the fraction of imports relative to local transmissions converged at the relatively similar values 0.35 and 0.21 when upsampling tips from ROW and Norway respectively. For Victoria, the corresponding numbers were 0.13 and 0.15, respectively (**Fig S7**). In contrast, the import estimates from the USA, similarly produced by either increasing the number of domestic genomes, or genomes from ROW, resulted in estimated import asymptotes at 0.06 and 0.14, respectively, a 2.3-fold difference (**Fig. S8**). For Europe, the discrepancy was even more pronounced, and the estimated fraction of imports did not converge for the upsampling of European genomes, while upsampling genomes from ROW yielded an asymptote at 0.2. We thus concluded that the sampling of Europe and the USA was insufficient for detailed quantitative analyses of imports in relation to local transmission.

Export asymptotes were also generated for all locations (**Figs. S9-S10**). The import and export asymptotes for Norway and Victoria, generated from point estimates from incrementally larger numbers of genomes from ROW, as described above, are presented in **Fig. 4**. For both locations, the distance between the point estimates of exports and the estimated asymptotes were markedly longer compared to the corresponding import analyses, indicating substantial uncertainty in the export estimates (Fig 4 B).

**Figure 4.**
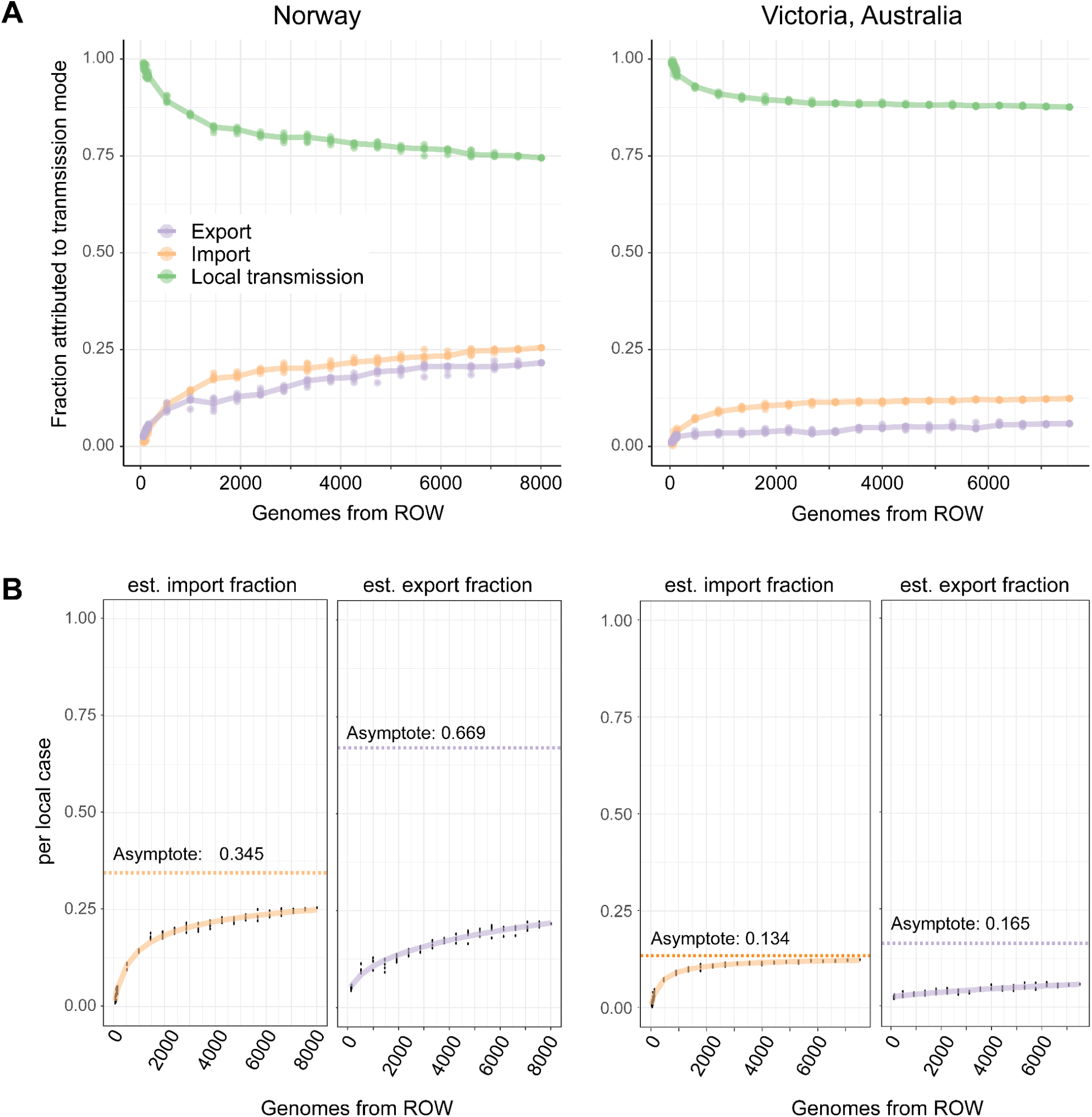
Assessing robustness of import and export estimates by upsampling. A) To assess the robustness of import and export estimates as a function of inclusion of samples from the rest of the world (ROW), the latter genomes were upsampled in a stepway manner from the dated phylogeny, followed by re-estimation and quantification of imports, exports and local transmissions. For each step, three random subsamples of ROW genomes were retained, and phylogeographic mapping estimated independently for each. B) Import and export fraction asymptotes were calculated to assess whether the estimates were stable and to estimate true fractions of imports and exports.

Despite the uncertainty surrounding export estimates for Norway and Victoria, the estimated frequency of exports per local transmission was consistently higher for Norway than Victoria (Figs. 5 and S9). This finding implies that a substantial fraction of cases in Norway are imports, but also frequently serves as a source of gonorrhea.

**Figure 5.**
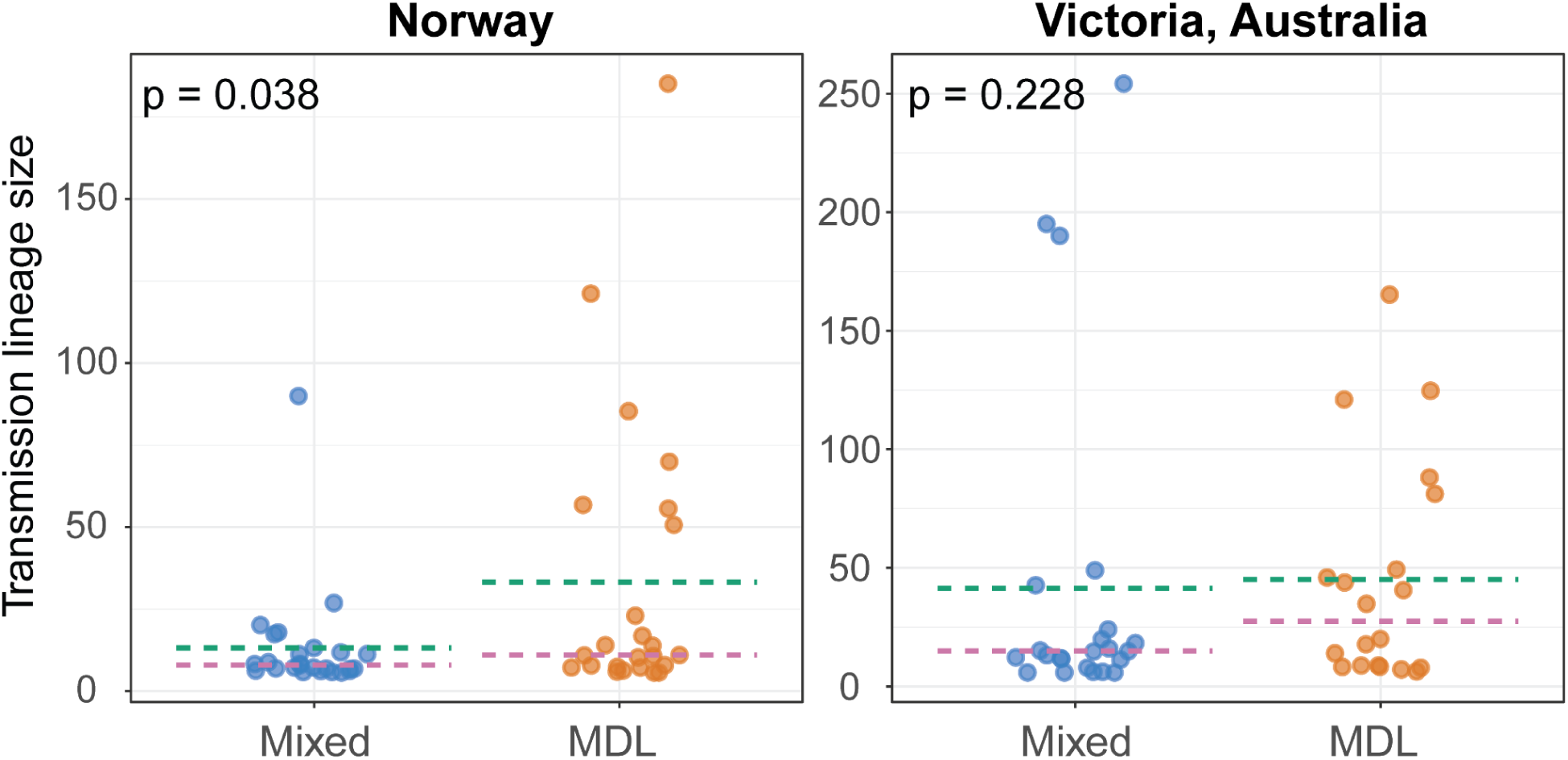
Transmission lineage sizes as a function of sex distribution. For each location, transmission lineages were annotated as male-dominated lineage (MDL) if they were in the top half of clusters in terms of male/female ratio and mixed otherwise. P-values are based on the DTS test (35). Blue: mixed lineages; orange: MDL lineages. The dotted lines indicate the median (magenta) and mean (green) transmission lineage size.

Next, we investigated the relationship between underlying putative sexual networks and transmission lineage size. Since we had no data directly indicative of sexual network structure, we used the observed fraction of men in the transmission lineage as proxy to assign lineages to network types. For each location (Norway and Victoria), transmission lineages containing more than 10 sequences were grouped into two groups based on the 50% percentile of the observed fraction of men in each transmission lineage. The 50% percentile was surprisingly stable across all four regions (85% both in Norway and in Europe excluding Norway, 87% in Australia, and 84% in the USA). As men were significantly overrepresented across all locations, lineages in the group with the lowest fraction of men were termed ‘mixed’ lineages, whereas those in the group with the highest fraction of men were termed male-dominated lineages (MDL). The ‘mixed’ group is thus expected to encompass lineages associated with both heterosexuals and men who have sex with men (MSM), whereas the MDL group is likely to be substantially enriched for gonococcal genomes from MSM.

In Norway, transmission lineages in the MDL group were significantly larger than those in the mixed group (Fig. 5), but no such association was found in Victoria. Moreover, we found a significant difference in the distribution of transmission lineage sizes between Norway and Victoria (DTS-test p=0.008). We estimated power-law distributions for lineage sizes in both regions. The scaling parameter (α) of the power-law distribution was 1.73 for Victoria and 2.03 for Norway, and the minimum values (*x_min_*) from which the power-law behavior started were 2 and 1, respectively (Kolmogorov-Smirnov test p-value > 0.99 for both regions). This suggests that the distribution for Victoria has a ’heavier’ tail, indicating a higher probability of observing very large lineages in Victoria compared to Norway.

## Discussion

### Global population dynamics and antibiotic resistance

Our temporal demographic analyses suggest that the global gonococcal effective population size has undergone nearly continuous expansion over the last two centuries (Fig. 2). However, between 2010 and 2018 (corresponding to the end of the study period), we identified a striking contraction in the effective population size. In apparent contradiction to this finding, the incidence of gonorrhea has increased steadily in Europe and the USA over the same period (22, 23, 36). When repeating the analyses on a subsampled genome collection, including only sequences from samples collected through unbiased sampling, the abrupt drop in effective population size from around 2010 was not reproduced, but still suggested a less dramatic decrease in effective population size in recent years (Fig. S2). Together, the demographic analyses of the full dataset (Fig. 2A) and the minimally biased dataset (Fig. S2) suggest that the effective population size of gonococci has expanded over two centuries, before likely plateauing or moderately declining in recent years.

A decrease in effective population does however not necessitate a decrease in incidence (25). Indeed, if a limited number of clades expand in a background of reduced genetic diversity, this could lead to a decrease in effective population size. A likely scenario is thus that the increase in gonorrhea incidence since ∼ 2010 has been driven by a limited number of successful clones. We recently showed that two clades of ST-1901 expanded as two distinct waves globally, and that the effective population size of each clade went into decline as new antibiotics, first cefixime (‘wave 1’), then ceftriaxone + azithromycin (‘wave 2’), were introduced (1). In addition, improved diagnostic tests and increased testing of extragenital sites were suggested to have contributed to the decline of ST-1901 between 2009/2010 and 2013/2014 (2). Improvements in both diagnostic sensitivity and treatment schemes are expected to introduce selective pressures, favoring increased transmissibility and/or shorter generation times, which could also introduce population bottlenecks.

Clade-specific demographic analyses suggested that the effective population size of three major clades carrying *penA* RS alleles have declined substantially since ∼ 2010, in contrast to the large clade carrying various mosaic versions of the *mtr* operon (Fig. 2). In addition, it is clear that *penA* RS and mosaic *mtr* alleles co-occurrence is exceedingly rare at the population level. In the context of gonorrhea therapy, reduced azithromycin susceptibility is expected to confer limited selective advantage on its own, as cephalosporins have been a cornerstone of recommended treatment regimes for two decades. We hypothesize that the switch from cefixime to increasingly higher doses of ceftriaxone nullified the selective advantage associated with carrying *penA* RS alleles, whereas the expansion of the *mtr* mosaic clade might partially be a bystander-effect. Macrolides are used to treat a broad range of infections, and as many gonorrhea infections go undiagnosed (37), exposure to macrolides in the absence of cephalosporins might be a relatively regular occurrence for gonococci. This could confer a selective advantage to strains harboring mosaic *mtr* alleles (38).

### Transmission lineage dynamics

Various methods are used to classify *N. gonorrhoeae* ‘types’. The multilocus sequence typing (MLST) scheme has been useful for surveillance, but lacks resolution, and is prone to homoplasies (30). Core-genome MLST represents an extension of the MLST concept, approaching the resolution of whole-genome SNP analyses (32). Here, we introduce the transmission lineage concept (39) to gonococci. The transmission lineage ‘typing’ adds information by defining lineages based on inferred migration history. We believe that inferring transmission lineage structure along with phylogenetic trees and traditional typing schemes is useful for understanding the epidemiology of gonorrhea in particular regions and the dynamics of intercontinental spread.

One of our motivations for describing transmission lineages is that they can potentially indicate source and sink locations in terms of global transmission dynamics. For example, if a certain location contains numerous ‘old’ transmission lineages, this is suggestive of sustained transmission in that location. Also, the transmission lineage framework allows for quantitative and comparative analyses of transmission networks between locations of interest.

Norway and Victoria were primarily characterized by younger transmission lineages, whereas Europe and the USA were characterized by larger and older lineages. This observation is to be expected as the choice of geographic levels directly affects the labeling of transmission lineages (for larger regions, a higher fraction of transmission events are identified as internal transmission events). Comparing Norway and Victoria, transmission lineages were significantly smaller in Norway, where imports were also found to be responsible for a higher fraction of new cases. We also found that male-dominated transmission lineages were significantly larger than mixed lineages in Norway, but not in Victoria (Fig. 5). Transmission lineage size can be shaped both by growth rate and age (time since import), and the interpretation of this finding is thus not straightforward. Furthermore, some lineages span both heterosexual and MSM networks, and even switch between different network-types over time, complicating simplistic analyses (14, 40). Taken together, our results indicate that the sex distribution within transmission networks plays a role in shaping the fate of transmission lineages.

Across all four regions of interest, transmission lineage sizes were compatible with being drawn from power law distributions. This could be related to the underlying structure of the sexual networks of the hosts, as the distribution of number of sexual partners is also known to be heavy-tailed and exhibit power law scaling (41, 42). However, as the same observation was made for SARS-CoV-2 transmission lineages (39), the power law distribution of transmission lineage sizes might reflect more general properties of infectious disease transmission networks. Such properties could either reflect the underlying transmission networks or pathogen properties, such as the dynamics of lineage survival and distribution of fitness differences arising through genetic variation generated over time.

### Import, export and local transmission dynamics

Regional differences in incidence are expected to have significant effects on relative rates of import and export. In the regions of study, the highest gonorrhea incidence is in the USA, with 206.5 reported cases per 100,000 inhabitants in 2020 (23). In Victoria, the yearly incidence over the three-year period 2017-2019 was 124 per 100,000 (43). In Norway, the reported incidence was much lower, at 31.3 per 100,000 in 2018, slightly higher than the European Economic Area average at 26.4 (36). In addition, travel frequency and destination of travel are important determinants of gonorrhea import and export rates. The lowest rates of imports and exports were inferred in Victoria, which might partially reflect relatively less dense sampling of nearby regions, but also the relative remoteness of Australia compared to Europe or North America.

We estimated both relative import and export rates of gonorrhea to and from Norway and Victoria and found much higher rates of both imports and exports to and from Norway. Together with the finding that transmission lineages were generally smaller in Norway, it seems clear that the gonorrhea epidemic in Norway is characterized by frequent imports generating shorter-lived transmission lineages compared to Victoria. The high rates of imports and export to and from Norway, combined with the large fraction of small and short-lived transmission lineages within the country, paint a picture of a national epidemiology with a high degree of cross-border interconnectedness. This likely reflects a combination of high rates of international travel and successful suppression of local transmission (30).

### Uncertainty in the assignment of transmission lineages and singletons

Here, we have extracted estimates of import, export and local transmissions from the distribution of transmission lineage and singletons obtained from a dated phylogeographic mapping. The resulting estimates of import, export and local transmissions are likely affected by multiple sources of uncertainty: collection bias, uncertainty in phylogenetic estimation, and dating estimation of the geographical states on the nodes. There exist approaches that could better accommodate this by including the uncertainty in an integrated probabilistic analysis (39). While it might be possible to account for various sources of uncertainty with a fully probabilistic modeling framework, this is not feasible when analyzing datasets of this size.

Collection bias limits the resolution of phylogeographic inferences (44) and will among other things impact the inferred size of transmission lineages. Furthermore, du Plessis *et. al.* (*39*) demonstrated that, for SARS-CoV-2, the time between importation and detection of transmission lineages is dependent on their sizes, with shorter lags for larger lineages. We expect this to be true for gonococci as well, so that our collection will be biased towards earlier observation of larger transmission lineages. If the collection does not represent the genetic diversity present in a geographical location, multiple imports from that location will be assigned to the same transmission lineage (Fig. S6). This can lead to an overestimation of transmission lineage size and hence the local transmission, and an underestimation of the number of imports using our approach.

Despite the uncertainties and limitations listed above, we expect large-scale transmission lineage dynamics to penetrate the uncertainty in large datasets. Our approach offers a pragmatic solution for rapidly estimating transmission lineages and the relative numbers of imports, exports and local transmission of an infectious disease.

### Comparison of quantitative estimates and self-reported travel histories

The availability of self-reported travel histories from the Norwegian Surveillance System for Communicable Diseases (22) enabled us to compare our inferences based solely on genomic and geographic data, with epidemiological data. Encouragingly, in the period 2016-2018, 25% of new cases in Norway represented new imports based on self-reporting, compared to 27% inferred herein, lending strength to our approach.

Aggregated self-reported data on whether patients had acquired gonorrhea abroad was available also for the Victoria dataset (14). In total, 5.6% of the patients in Victoria reported having had ‘overseas sex’ within a time-frame compatible with having acquired the infection abroad. This number is significantly lower than our estimates at 12.7% for the state of Victoria. One obvious explanation for the discrepancy is that we treat the state of Victoria, rather than Australia, as the focal location in our analyses. Also, despite dense sampling of Victoria, the rest of Australia was hardly sampled at all. Hence, any strain entering Victoria from some other part of Australia, and which is more closely related to a sampled genome in the rest of the world than to previously observed samples in Victoria, will be identified as an import event. By comparing relative import estimates from our quantitative analyses (12.7%) with those based on self-reporting (5.6%), we can estimate that a little over half of new infections imported to Victoria came from other Australian states and a little less than half came from abroad. These estimates should however be treated with caution, as there is no easy way to assess their accuracy.

## Conclusions

By combining computationally efficient methods for masking recombination, generating temporal phylogenies, reconstructing and summarizing phylogeographic histories, we were able to detect and summarize region-specific gonorrhea transmission dynamics. We believe the approach presented herein represents an attractive framework for the exploration of intercontinental pathogen dynamics beyond gonorrhea.

We show that, given sufficient sampling, the relative rate of importation of *N. gonorrhoeae* can be quantified with high accuracy. In addition, a novelty of the current work is that we explicitly highlight infectious disease export as a concept to consider. Public health authorities clearly recognize imports as a source of new infections, and routinely give advice with the aim of reducing imports and onward transmission. However, any import of a pathogen to a new location can also be regarded as an export from some other location. We show that relative rates of gonorrhea export can be quantified, but require dense sampling of both origin and export locations, in contrast with the detection of imports, which can be performed using a more limited number of extraneous sequences, or even using only samples from the region of interest (45). Despite the uncertainty surrounding the export estimates, our analyses suggest that some regions export gonorrhea at higher rates than others. Export estimates can be used to create awareness both nationally and internationally, and to guide targeted interventions both at the source and in the receiving end, via the identification of high-risk areas

## Methods

### Data collection

A total of 9,732 genomes were included in the study. Collection date was known for all samples, and gender information was successfully retrieved for 6796 (70%) of the patients, including all samples in the Norway, USA and Victoria datasets described below. The collection was built by selecting all gonococcal genomes (n=1,842) available at PathogenWatch (pathogen.watch) as of November 5th 2019. All of these had been sampled in 2013 or earlier. To obtain better coverage of more recent samples, all genomes sampled in 2014 or later were retrieved from the PubMLST server (n=805) (46) as of September 23rd 2019. In addition, selected genomes were subsequently added to improve spatial and temporal coverage, specifically 121 genomes from Vietnam 2016 (47) and historical genomes from Denmark (26); USA collection: 2,144 genomes from USA 2016-2018 (PRJNA317462, accessed on December 16th 2019); Victoria collection: 2,184 genomes collected in the state of Victoria, Australia throughout 2017 (14); Norway collection: 1658 genomes sequenced at the Norwegian Institute of Public Health, representing all culture-positive and successfully sequenced cases in Norway 2016-2018 (PRJEB32435); 318 genomes from the Public Health Service of Amsterdam, The Netherlands, 2014-2019 (48); 223 genomes from the ’Gentamicin for the treatment of gonorrhoea’ study (27) collected in the UK 2014-2016; 437 genomes from China 2011-2013 (PRJNA431691) and 2017-2018 (PRJNA560592).

### Data availability

All the metadata for the project can be accessed on Zenodo at https://doi.org/10.5281/zenodo.8055415.

### Genome assembly

Assemblies in fasta format were downloaded directly from PubMLST and PathogenWatch. Sequence data from other sources were assembled as follows: FastQ reads were trimmed using Trimmomatic v.0.38 (https://github.com/timflutre/trimmomatic) and assembled using Spades v.13.0 (49) using ‘careful’ mode and automatic coverage cutoff. The assemblies were filtered by size (excluding files > 2.3 mb or < 2.0 mb) and all contigs shorter than 500 nt and/or with a Spades coverage < 2 removed using ‘filter_bad_contigs.py’ (https://github.com/AdmiralenOla/fhiscripts/blob/master/filter_bad_contigs.sh).

### Filtering of highly recombinant regions of the genome

As standard approaches for identifying recombinant regions (19, 20) are unable to handle datasets containing ∼10,000 bacterial genomes, we employed a novel approach to mask the most recombination-heavy regions prior to phylogenetic reconstruction.

First, five random subsets containing 500 genomes were drawn from the total collection of 9,732 genomes. Whole-genome alignments were generated by combining Parsnp (50) with ‘parsnp2fasta’ (https://github.com/krepsen/parsnp2fasta) as described (1). Subsequently, recombination events were identified using Gubbins (19) on each subset. We subsequently identified the most recombined regions across the subsets, defined as belonging to either the 10% of sites in terms of (**1**) number of unique recombination events, or (**2**) taxa affected by recombination. We then masked all sites identified as most recombined according to the two schemes described above, resulting in a total of 12% of all sites in the alignment. We found that 64-72% of all recombinant sites in each of the five subsets were successfully masked when applying this cross-subset filtering step. The final list of masked sites included 73 regions covering a total of 262,597 sites out of a total number of 2,153,922 sites in the FA1090 reference genome (Table S2 and Fig S11).

To assess the influence of masking pre-defined recombinant regions, we contrasted dated trees produced with and without masking of the 73 regions. For each tree, root position analysis underscored a single region in the tree with a superior correlation between root-to-tip distance and collection date (Fig S12). The temporal signal was similar before and after masking highly recombinant regions (R2 = 0.14 vs R2 = 0.13, see Fig. S12).

Using LSD2 we built dated trees (51) and estimated substitution rates (see the *Phylogenetic tree estimation and dating* below). Masking recombination led to a decrease in the estimated substitution rate, from 1.33x10^-5^ (95% CI: 1.29x10^-5^ - 1.37x10^-5^) to 7.41x10^-6^ (95% CI: 7.16x10^-6^ - 7.75x10^-6^) substitutions/site/year. This corresponds to a shift from 28.83 (95% CI: 27.95 - 29.60) to 14.03 (95% CI: 13.54 - 14.66) substitutions/genome/year. The rate estimated from the recombination-masked phylogenetic analysis is at the higher end of recent genome-wide phylodynamic-inferred estimates (1, 26, 30). Moreover, we explored differences in dating close and more distant ancestral relationships by examining the variation in the times to the most recent ancestors (TMRCAs) of leaves and their 1, 2, 5, 10, 20, and 50 closest neighbors in the trees (see Supplementary Methods). The analysis revealed that the rooted, recombination-masked tree estimated younger TMRCAs for most groups of leaves (Fig S13), and that the difference between the TMRCAs in the trees narrowed towards the present (Fig S14). Taken together, our results illustrate a discernible, albeit modest, effect of masking the most recombinogenic regions, with the most pronounced effect being reduced and likely more realistic mutation rate estimates.

### Generation of recombination-corrected whole-genome alignment

Genomes were aligned to the FA1090 reference (NC_002946.2) genome using NUCmer v.3.1 (52) with the ‘maxmatch’ option. A whole-genome alignment was created from the resulting SNP files using the script ‘WGA_from_mummer.py’ (https://github.com/AdmiralenOla/fhiscripts/blob/master/WGA_from_mummer.py). For each SNP file, the script generates a new fasta from the reference genome, with identified SNPs replacing the corresponding position in the reference. SNP-sites (53) was used to output a whole-genome SNP alignment containing only variable sites. The above approach was run both on the regular FA1090 reference genome and on a version of FA1090 where the 73 highly recombined regions had been replaced by Ns (hence retaining positional information, but ignoring and SNPs in these regions). The final whole-genome SNP alignment generated from the recombination-masked FA1090 reference included a total of 168,013 variable positions, and was used for phylogenetic reconstruction below.

### Characterization of resistance alleles

To characterize alleles of interest associated with reduced susceptibility to cephalosporins and azithromycin, we followed the approach outlined in Osnes et al (1). Briefly, whole genome assemblies were aligned against gene-specific databases downloaded from pubMLST (46) using BLAST (54). This was done for the following genes/segments: *penA* (NEIS1753), *mtrR* (‘mtrR – includes promoter region) and *23S* (NG_23S). We retained up to four hits from the 23S BLAST hits as multiple copies are generally present in the chromosome. All isolates, for which at least one allele were identified as bearing a A2059G or C2611G mutation, were counted as positive hits. For *penA* alleles, we used the NG-STAR nomenclature (55). In addition, to identify and characterize mosaic *mtrCDE* alleles associated with azithromycin resistance (13, 14), the region encompassing the three-gene operon in the FA1090 reference genome (positions 1321639-1327548) were extracted from the whole-genome alignment. The extracted *mtrCD*E alignment was subsequently used as input for analysis in fastGEAR (29) specifying default settings.

We assigned *penA* alleles as mosaic or non-mosaic based on NG-STAR classifications. In addition, we used a semi-manual approach to assign penA alleles as ‘high’ or ‘low’ cephalosporin MIC alleles independent of the presence of mosaicism. Each allele was assigned as either ‘high’ or ‘low’ based on 1) NG-STAR annotations, 2) published data, or 3) available MIC measurements at Norwegian Institute of Public Health. The cut-off for being placed in the ‘high’ category was that the allele was associated with an MIC of ≥ 0.064 for cefixime and/or ceftriaxone (30). The applied method is pragmatic, and it should be noted that the evidence, that is, the number of measurements, is limited for some alleles.

### Phylogenetic tree estimation and dating

We estimated phylogenetic trees using FastTree v. 2.1.10 (56), using the Jukes-Cantor substitution model. The tree was rooted by selecting the midpoint of the edge that yielded the highest correlation between root-to-tip distance and collection date. Using LSD2 v. 2.4.4 (51), we generated a time-scaled phylogeny derived from the genetic distances. Within the tree, two of the clades corresponded to those for which dated trees had been estimated in prior studies. Consequently, we calibrated the ancestral nodes of ST-7827 (30) and ST-1901 (1) to ensure our dating was in agreement with these previous estimates.

### Estimating demographic histories

Effective population size through time was reconstructed using skygrowth v.0.3.1 (21) with default settings. Demographic histories were reconstructed independently from the full dated tree (9,732 genomes) and based only on the dated tree built from unbiased sequence collections (4184 genomes from Norway, Australia (14), Denmark (26) and the UK (27)).

### Phylogeographic mapping

We were interested in describing the frequency of local transmission versus importation for different geographical regions. The collection contained a high sampling density for Norway (n=1724), Australia (n=2203), and the USA (n=2372), compared to 3434 sequences from all other locations. We therefore conducted three separate geographical analyses for Norway, Australia and the USA, with in each case two categories: the respective location and the rest of the world (RoW). We mapped the geographical locations on the phylogenetic tree using maximum likelihood as implemented in the ace function in the ape v.5.6-1 (33) R-package with the default settings. We used the all-rates-different (ARD) model to allow for unequal transition rates between the geographical regions.

### Estimating local transmission clusters

Using the mapped geographical locations, we defined local transmission lineages using LineageHomology (34, 57). LineageHomology defines transmission lineages in the same fashion as (39), but here we have used it in conjunction with maximum likelihood reconstruction of the geographical locations. The internal nodes of the tree are labeled with the geographical location that is estimated with more than 50% probability (there is always such a location since we only consider two states in each phylogeographic analysis). Sequences are assigned to the same local transmission lineage if they are mapped to the same geographical locations without any estimated geographical transitions on the path that connects these sequences in the phylogeny. If single sequences are mapped to a different location than the neighboring sequences, these are labeled as singletons.

### Transmission lineage statistics

To describe the differences in the properties of the transmission lineages in the different locations we extracted summary statistics from the transmission clusters estimated using *LineageHomology*. For each location, we calculated the number of cases attributed to its ten largest transmission lineages. Additionally, we determined the number of cases that could be explained by singletons.

We observed significant variation in the size of transmission lineages with a heavy tail towards very large sizes. Therefore, we utilized the igraph R-package (58) to fit a power law distribution to the counts of lineage sizes in each location. This approach assumes that the transmission lineage size (x) is proportional to a power law: *x^−α^*, where the power law distribution describes the data after a minimum value (x_min_) has been reached. To assess the compatibility of the data with the estimated power law distributions, we calculated the p-value using the Kolmogorov-Smirnov test.

To compare the distribution of transmission lineage sizes, we employed the DTS-test implemented in the twosamples R-package (59).

### Deriving import, export and local transmission from the estimated transmission lineages

For each of the locations, we used LineageHomology to describe importation events (IEs), local transmission events (LTEs) and exportation events (EEs) over time. Importantly LineageHomology does not consider unobserved sequences. The IEs, LTEs, and EEs are based on the phylogeographic mapping of the observed sequences. LTEs are assumed to occur when there are branching events within a transmission lineage. The time of the LTE is set to the estimated time of the node that defines the branching event, which is extracted from the apriori estimated timetree. IEs are assumed to occur at the root branch of all transmission lineages (as these are defined based on geographical transitions). The time of the IE is set to the midpoint of the branch that leads to the root node of the transmission lineage. EEs are assumed to occur when there is a branching event and a downstream node is estimated to be in a different geographical location. The time of the EE is assumed to be at the midpoint of the branch leading to the other location. Note that an IE and EE can represent the same geographical transition, but are defined as an IE or EE based on the focal geographical location (Fig. S5). We obtained uncertainty estimates by using LineageHomology to draw the geographical states of the nodes 100 times based on maximum likelihood probabilities of the geographical states, and then redefining transmission lineages based on the draws. Hence each replicate yields a new stochastic draw of transmission lineages, IEs, LTEs, and EEs, which allows us to represent how the uncertainty in the phylogeographic mapping translates to uncertainty in the timing of IEs, LTEs, and EEs.

### Effective population size estimation and transmission lineage growth rate analysis

We used skygrowth v.0.3.1 (21) with default settings to estimate the effective population size in the full tree and clades of interest. The clades of interest were distinguished based on *mtr* lineage assignment by FastGear and assigned *penA* alleles and types. For each of the clades, we visually assessed the position of the root node and included all downstream branches in the corresponding skygrowth run. Additionally, we ran skygrowth on all transmission lineages that contained more than 10 sequences and extracted the effective population size and growth rate in the period 2010 - 2017 to describe the recent growth and size of the underlying pathogen populations of the transmission lineages.

### Assessing the effect of sex distribution on transmission lineage sizes

We sought to determine if male-dominated transmission lineages exhibit larger sizes compared to those with a higher proportion of women. We calculated the male-to-female ratio for each transmission lineage and ranked them according to the relative percentage of men. A breakpoint was established at the 50th percentile for each geographical region: Norway, Australia, Europe, and the USA. Lineages above this breakpoint were designated as male-dominated lineages (MDL), while those below were labeled mixed lineages, signifying a higher proportion of women. The DTS test (35) was employed to compare the distribution of transmission lineage sizes between MDL and mixed groups, and to calculate the p-value of the observed difference.

### Assessing the suitability of sample collections for quantifying imports, exports, and local transmission

We used an “upsampling” approach to evaluate the phylogeographic signal of sample collections for accurate relative quantification of imports, exports, and local transmission in different geographical regions. We performed the analysis for Norway, Victoria, Europe, and the USA. We upsampled tips in the dated phylogeny from the focal location and the Rest of the World (ROW) independently. For example, in the analysis focusing on Norway, ROW consisted of 8007 genomes in total, and in the upsampling subsets we included 57, 77, 97, 117, … , 6605, 7072, 7540, 8007. For each subset, the following steps five times were replicated, 1) randomly including x genomes from the upsampling location from the dated phylogeny - in practice we removed all other genomes than those selected for inclusion 2) estimate geographic location of the nodes using maximum likelihood in ace 3) assigned transmission lineages to the mapping using LineageHomology 4) calculated import, export and local transmission as before. This procedure was performed both for the focal location and for the external location ROW to estimate convergence of import, export, and local transmission estimates.

We noted that the imports and exports increases from zero and dampens towards a value as we approach the full collection of genomes from ROW. The local transmission estimate starts at 100 percent and decreases in a dampening fashion as we include more genomes from ROW, because more genomes are identified as imports.

To determine if the estimates were close to convergence, indicating a sufficient number of samples from the focal locations for accurate geographic reconstructions, we used nonlinear least squares various to fit the asymptotic functions in the drc R-package (60). We selected the model with the lowest residual mean squared error (RMSE) among the asymptotic functions and reported the asymptotes as the estimated parameter of this model. We considered the phylogeographic signal sufficient if the asymptote was similar when upsampling of the focal and upsampling of ROW yielded asymptotes at similar points.

## Notes

https://magnusnosnes.github.io/10000_Ngon_genomes/

### Competing Interest Statement

The authors have declared no competing interest.

## References

1. M. N. Osnes, et al., Antibiotic treatment regimes as a driver of the global population dynamics of a major gonorrhea lineage. Mol. Biol. Evol. (2020) https://doi.org/10.1093/molbev/msaa282 (November 24, 2020).

2. S. R. Harris, et al., Public health surveillance of multidrug-resistant clones of Neisseria gonorrhoeae in Europe: a genomic survey. Lancet Infect. Dis. 18, 758–768 (2018).

3. L. Sánchez-Busó, et al., The impact of antimicrobials on gonococcal evolution. Nat Microbiol 4, 1941–1950 (2019).

4. C. Bignell, M. Unemo, 2012 European guideline on the diagnosis and treatment of gonorrhoea in adults. Int. J. STD AIDS 24 (2013).

5. Update to CDC’s Sexually Transmitted Diseases Treatment Guidelines, 2010: Oral Cephalosporins No Longer a Recommended Treatment for Gonococcal Infections (2012) (September 2, 2022).

6. H. Fifer, J. Saunders, S. Soni, S. T. Sadiq, M. FitzGerald, 2018 UK national guideline for the management of infection with Neisseria gonorrhoeae. Int. J. STD AIDS 31, 4–15 (2020).

7. S. St Cyr, et al., Update to CDC’s Treatment Guidelines for Gonococcal Infection, 2020. MMWR Morb. Mortal. Wkly. Rep. 69, 1911–1916 (2020).

8. S. Ameyama, et al., Mosaic-like structure of penicillin-binding protein 2 Gene (penA) in clinical isolates of Neisseria gonorrhoeae with reduced susceptibility to cefixime. Antimicrob. Agents Chemother. 46, 3744–3749 (2002).

9. B. G. Spratt, L. D. Bowler, Q. Y. Zhang, J. Zhou, J. M. Smith, Role of interspecies transfer of chromosomal genes in the evolution of penicillin resistance in pathogenic and commensal Neisseria species. J. Mol. Evol. 34, 115–125 (1992).

10. K. C. Ma, et al., Increased power from bacterial genome-wide association conditional on known effects identifies Neisseria gonorrhoeae macrolide resistance mutations in the 50S ribosomal protein L4. Cold Spring Harbor Laboratory, 2020.03.24.006650 (2020).

11. Y. H. Grad, et al., Genomic Epidemiology of Gonococcal Resistance to Extended-Spectrum Cephalosporins, Macrolides, and Fluoroquinolones in the United States, 2000-2013. J. Infect. Dis. 214, 1579–1587 (2016).

12. L. Zarantonelli, G. Borthagaray, E. H. Lee, W. M. Shafer, Decreased azithromycin susceptibility of Neisseria gonorrhoeae due to mtrR mutations. Antimicrob. Agents Chemother. 43, 2468–2472 (1999).

13. C. B. Wadsworth, B. J. Arnold, M. R. A. Sater, Y. H. Grad, Azithromycin Resistance through Interspecific Acquisition of an Epistasis-Dependent Efflux Pump Component and Transcriptional Regulator in Neisseria gonorrhoeae. MBio 9 (2018).

14. D. A. Williamson, et al., Bridging of Neisseria gonorrhoeae lineages across sexual networks in the HIV pre-exposure prophylaxis era. Nat. Commun. 10, 3988 (2019).

15. L. Sánchez-Busó, et al., A community-driven resource for genomic epidemiology and antimicrobial resistance prediction of Neisseria gonorrhoeae at Pathogenwatch. Genome Med. 13, 61 (2021).

16. D. De Silva, et al., Whole-genome sequencing to determine transmission of Neisseria gonorrhoeae: an observational study. Lancet Infect. Dis. 16, 1295–1303 (2016).

17. K. Town, et al., Genomic and Phenotypic Variability in Neisseria gonorrhoeae Antimicrobial Susceptibility, England. Emerg. Infect. Dis. 26, 505–515 (2020).

18. X. Didelot, J. Parkhill, A scalable analytical approach from bacterial genomes to epidemiology. Philos. Trans. R. Soc. Lond. B Biol. Sci. 377, 20210246 (2022).

19. N. J. Croucher, et al., Rapid phylogenetic analysis of large samples of recombinant bacterial whole genome sequences using Gubbins. Nucleic Acids Res. 43, e15 (2015).

20. X. Didelot, D. J. Wilson, ClonalFrameML: efficient inference of recombination in whole bacterial genomes. PLoS Comput. Biol. 11, e1004041 (2015).

21. E. M. Volz, X. Didelot, Modeling the Growth and Decline of Pathogen Effective Population Size Provides Insight into Epidemic Dynamics and Drivers of Antimicrobial Resistance. Syst. Biol. 67, 719–728 (2018).

22. MSIS, the Norwegian Surveillance System for Communicable Diseases. MSIS-statistikk (October 1, 2021).

23. Centers for Disease Control and Prevention, “Preliminary 2021 STD Surveillance Data” (CDC, 2021).

24. Centers for Disease Control and Prevention, “Gonococcal Isolate Surveillance Project (GISP) and Enhanced GISP (eGISP) Protocol” (CDC, 2020).

25. B. Charlesworth, Effective population size and patterns of molecular evolution and variation. Nat. Rev. Genet. 10, 195–205 (2009).

26. D. Golparian, et al., Genomic evolution of Neisseria gonorrhoeae since the preantibiotic era (1928-2013): antimicrobial use/misuse selects for resistance and drives evolution. BMC Genomics 21, 116 (2020).

27. J. D. C. Ross, et al., Gentamicin compared with ceftriaxone for the treatment of gonorrhoea (G-ToG): a randomised non-inferiority trial. Lancet 393, 2511–2520 (2019).

28. K. Alfsnes, et al., Genomic epidemiology and population structure of Neisseria gonorrhoeae in Norway, 2016-2017. Microb Genom (2020) https://doi.org/10.1099/mgen.0.000359.

29. R. Mostowy, et al., Efficient Inference of Recent and Ancestral Recombination within Bacterial Populations. Mol. Biol. Evol. 34, 1167–1182 (2017).

30. M. N. Osnes, et al., Sudden emergence of a Neisseria gonorrhoeae clade with reduced susceptibility to extended-spectrum cephalosporins, Norway. Microb Genom (2020) https://doi.org/10.1099/mgen.0.000480.

31. K. Yahara, et al., Emergence and evolution of antimicrobial resistance genes and mutations in Neisseria gonorrhoeae. Genome Med. 13, 51 (2021).

32. O. B. Harrison, et al., Neisseria gonorrhoeae Population Genomics: Use of the Gonococcal Core Genome to Improve Surveillance of Antimicrobial Resistance. J. Infect. Dis. (2020) https://doi.org/10.1093/infdis/jiaa002.

33. E. Paradis, K. Schliep, ape 5.0: an environment for modern phylogenetics and evolutionary analyses in R. Bioinformatics 35, 526–528 (2019).

34. M. N. Osnes, et al., The Impact of Global Lineage Dynamics, Border Restrictions and Emergence of the B.1.1.7 Lineage on the SARS-CoV-2 Epidemic in Norway. Virus Evol (2021) https://doi.org/10.1093/ve/veab086 (September 29, 2021).

35. C. Dowd, A New ECDF Two-Sample Test Statistic. arXiv [stat.ME] (2020).

36. European Centre for Disease Prevention and Control, “Gonorrhoea - Annual Epidemiological Report for 2018” (ECDC, 2020).

37. R. D. Assaf, N. J. Cunningham, P. C. Adamson, J. T. Jann, R. K. Bolan, High proportions of rectal and pharyngeal chlamydia and gonorrhoea cases among cisgender men are missed using current CDC screening recommendations. Sex. Transm. Infect. 98, 586–591 (2022).

38. S. W. Olesen, et al., Azithromycin Susceptibility Among Neisseria gonorrhoeae Isolates and Seasonal Macrolide Use. J. Infect. Dis. 219, 619–623 (2019).

39. L. du Plessis, et al., Establishment and lineage dynamics of the SARS-CoV-2 epidemic in the UK. Science 371, 708–712 (2021).

40. K. Town, et al., Phylogenomic analysis of Neisseria gonorrhoeae transmission to assess sexual mixing and HIV transmission risk in England: a cross-sectional, observational, whole-genome sequencing study. Lancet Infect. Dis. 20, 478–486 (2020).

41. F. Liljeros, C. R. Edling, L. A. Amaral, H. E. Stanley, Y. Aberg, The web of human sexual contacts. Nature 411, 907–908 (2001).

42. B. F. de Blasio, A. Svensson, F. Liljeros, Preferential attachment in sexual networks. Proc. Natl. Acad. Sci. U. S. A. 104, 10762–10767 (2007).

43. E. P. F. Chow, C. K. Fairley, D. A. Williamson, M. Y. Chen, Spatial mapping of gonorrhoea notifications by sexual practice in Victoria, Australia, 2017-2019. Aust. N. Z. J. Public Health 45, 672–674 (2021).

44. N. De Maio, C.-H. Wu, K. M. O’Reilly, D. Wilson, New Routes to Phylogeography: A Bayesian Structured Coalescent Approximation. PLoS Genet. 11, e1005421 (2015).

45. X. Didelot, D. Helekal, M. Kendall, P. Ribeca, Distinguishing imported cases from locally acquired cases within a geographically limited genomic sample of an infectious disease. bioRxiv, 2022.07.15.500228 (2022).

46. K. A. Jolley, J. E. Bray, M. C. J. Maiden, Open-access bacterial population genomics: BIGSdb software, the PubMLST.org website and their applications. Wellcome Open Res 3, 124 (2018).

47. P. T. Lan, et al., Genomic analysis and antimicrobial resistance of Neisseria gonorrhoeae isolates from Vietnam in 2011 and 2015-16. J. Antimicrob. Chemother. 75, 1432–1438 (2020).

48. J. de Korne-Elenbaas, S. M. Bruisten, H. J. C. de Vries, A. P. Van Dam, Emergence of a Neisseria gonorrhoeae clone with reduced cephalosporin susceptibility between 2014 and 2019 in Amsterdam, The Netherlands, revealed by genomic population analysis. J. Antimicrob. Chemother. 76, 1759–1768 (2021).

49. A. Bankevich, et al., SPAdes: a new genome assembly algorithm and its applications to single-cell sequencing. J. Comput. Biol. 19, 455–477 (2012).

50. T. J. Treangen, B. D. Ondov, S. Koren, A. M. Phillippy, The Harvest suite for rapid core-genome alignment and visualization of thousands of intraspecific microbial genomes. Genome Biol. 15, 524 (2014).

51. T.-H. To, M. Jung, S. Lycett, O. Gascuel, Fast Dating Using Least-Squares Criteria and Algorithms. Syst. Biol. 65, 82–97 (2016).

52. S. Kurtz, et al., Versatile and open software for comparing large genomes. Genome Biol. 5, R12 (2004).

53. A. J. Page, et al., SNP-sites: rapid efficient extraction of SNPs from multi-FASTA alignments. Microb Genom 2, e000056 (2016).

54. S. F. Altschul, W. Gish, W. Miller, E. W. Myers, D. J. Lipman, Basic local alignment search tool. J. Mol. Biol. 215, 403–410 (1990).

55. W. Demczuk, S. Sidhu, M. Unemo, Neisseria gonorrhoeae sequence typing for antimicrobial resistance, a novel antimicrobial resistance multilocus typing scheme for tracking global dissemination of N …. Journal of clinical (2017).

56. M. N. Price, P. S. Dehal, A. P. Arkin, FastTree 2--approximately maximum-likelihood trees for large alignments. PLoS One 5, e9490 (2010).

57. M. N. Osnes, LineageHomology. GitHub repository (2021).

58. G. Csardi, T. Nepusz, The igraph software package for complex network research. InterJournal Complex Systems, 1695 (2006).

59. C. Dowd, twosamples: Fast Permutation Based Two Sample Tests (2023).

60. C. Ritz, F. Baty, J. C. Streibig, D. Gerhard, Dose-Response Analysis Using R. PLOS ONE 10 (2015).

