## supplementary material for "The inter-continental population dynamics of *Neisseria gonorrhoeae*"

### Table of contents

**SUPPLEMENTARY METHODS**

The effect of masking recombination on time to the most recent common ancestor 1

**SUPPLEMENTARY TABLES** 2

Azithromycin MIC by mtr FastGear lineage 2

List of highly recombined regions 2

Description of downsampled datasets 3

**SUPPLEMENTARY FIGURES** 4

Collection dates of the genomes 4

Effective population size in downsampled and the full dataset 5

23S and mtr alleles mapped to the dated phylogeny 6

Azithromycin MICs stratified by mtr lineage as identified by FastGear 6

How LineageHomology derives import, export and local transmission 7

The impact of collection bias on the estimates from LineageHomology 8

Import and export fraction asymptote estimates 9-12

Details on recombination filtering 13

Root position and temporal signal analysis 14

The estimated time to the most recent common ancestor (TMRCA) for recombination-masked and unmasked trees. 15-16

**References** 17

**Investigating the effect of masking recombination on time to the most recent common ancestor (TMRCA)**

To examine the influence of our recombination masking procedure on temporal estimates, we analyzed differences in TMRCA between the ancestors of groups of tips drawn from two distinct trees. These trees were independently estimated and rooted, with an aim to optimize the correlation between root-to-tip distance and time. Consequently, the tree topologies differed from each other.

For comparison, we selected groups of leaves in varying sizes from the trees, identified the most recent common ancestor (MRCA) in each tree, and compared them. Leaf groups were defined by traversing the tip labels of the recombination-masked tree. For each leaf, groups were determined by choosing the 1, 2, 5, 10, 20, and 50 closest neighbors of that leaf in the tree. We then established the TMRCA of these leaves in both the recombination-masked and unmasked trees. After this, the selected leaf and its neighbors were removed from the leaf traversal list and the process was repeated until the leaf list was exhausted.

We graphed the TMRCAs against each other and compared the points to a line with slope one and intercept zero. Next, we plotted the deviations over the entire timeline (Fig SX), specifically between the years 1980 and 2018, to distinctly highlight differences in more recent events. To emphasize areas of highest point density, we calculated rolling percentiles (97.5, 75, 62.5, 37.5, 25, 2.5) at intervals over a 15-year period.

The plots indicate that for the tree with masked recombination, most TMRCAs are estimated to be younger. This suggests that the masked tree has relatively younger dates, and consequently. This could also be a result of topological differences as the trees were estimated and rooted independently. The variation in this difference increases towards older TMRCAs and lessens towards the present.

| **Table S1. Azithromycin MIC by mtr FastGear lineage. The table summarizes MICs for 970 isolates (of a total of 1517 isolates with measured MIC), after removing isolates with known resistance mutations in rrl (23S) and the mtrR promoter** | | | | | |
|---|---|---|---|---|---|
| **Lineage** | **mean_MIC** | **max_MIC** | **min_MIC** | **median_MIC** | **stdev_MIC** |
| 1 | 1.047 | 2 | 0.094 | 1.047 | 1.3477455 |
| 2 | 0.758 | 1 | 0.032 | 1 | 0.484 |
| 3 | 2.0028235 | 4 | 0.032 | 2 | 0.8094302 |
| 4 | 0.5515873 | 2 | 0.25 | 0.5 | 0.237313 |
| 5 | 0.1945659 | 1 | 0.016 | 0.125 | 0.1503176 |

| Table S2. Final list of highly recombined regions identified and filtered out (relative to NC_002946.2) | |
|---|---|
| **start position** | **end position** |
| 57278 | 58417 |
| 67419 | 74005 |
| 96456 | 97760 |
| 125033 | 144188 |
| 163398 | 165893 |
| 235815 | 238644 |
| 273989 | 276891 |
| 293738 | 297770 |
| 375895 | 376501 |
| 384779 | 387348 |
| 402439 | 402483 |
| 404613 | 405770 |
| 426912 | 430622 |
| 433560 | 437463 |
| 441532 | 456131 |
| 503641 | 510910 |
| 562315 | 562856 |
| 568481 | 572821 |
| 632801 | 634503 |
| 634612 | 634612 |
| 674131 | 678153 |
| 690885 | 696898 |
| 791739 | 792462 |
| 842955 | 843617 |
| 866238 | 871683 |
| 914409 | 928804 |
| 933968 | 937710 |
| 938046 | 938152 |
| 941629 | 941693 |
| 955776 | 962714 |
| 1003169 | 1003169 |
| 1003192 | 1007731 |
| 1038573 | 1041265 |
| 1155425 | 1155476 |
| 1155488 | 1155992 |
| 1156124 | 1156331 |
| 1156469 | 1156943 |
| 1230389 | 1230408 |
| 1235028 | 1235028 |
| 1235590 | 1235671 |
| 1327884 | 1335862 |

| | |
|---|---|
| 1401930 | 1410159 |
| 1418122 | 1433375 |
| 1456936 | 1462913 |
| 1472612 | 1478037 |
| 1494781 | 1496191 |
| 1530828 | 1537474 |
| 1568857 | 1569711 |
| 1570150 | 1570150 |
| 1570292 | 1570322 |
| 1573276 | 1573458 |
| 1668146 | 1670152 |
| 1697106 | 1705252 |
| 1735388 | 1736768 |
| 1761425 | 1761953 |
| 1776205 | 1776214 |
| 1776564 | 1776564 |
| 1777254 | 1777760 |
| 1783440 | 1787983 |
| 1829413 | 1836107 |
| 1843436 | 1843922 |
| 1894411 | 1898550 |
| 1999068 | 2006542 |
| 2017704 | 2029336 |
| 2029936 | 2029936 |
| 2030148 | 2030250 |
| 2030682 | 2030779 |
| 2030795 | 2035716 |
| 2050718 | 2056949 |
| 2065589 | 2074267 |
| 2114622 | 2115855 |
| 2128064 | 2131435 |
| 2137280 | 2144085 |

#### SUPPLEMENTARY FIGURES

**A**

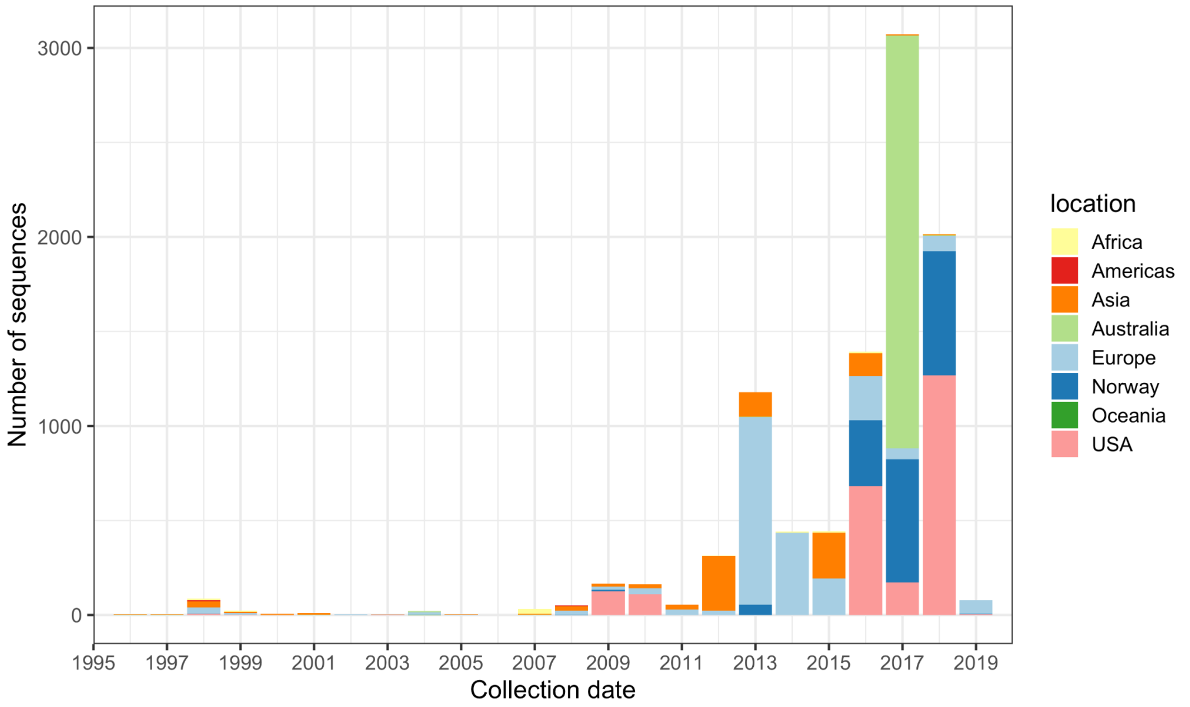


**B**

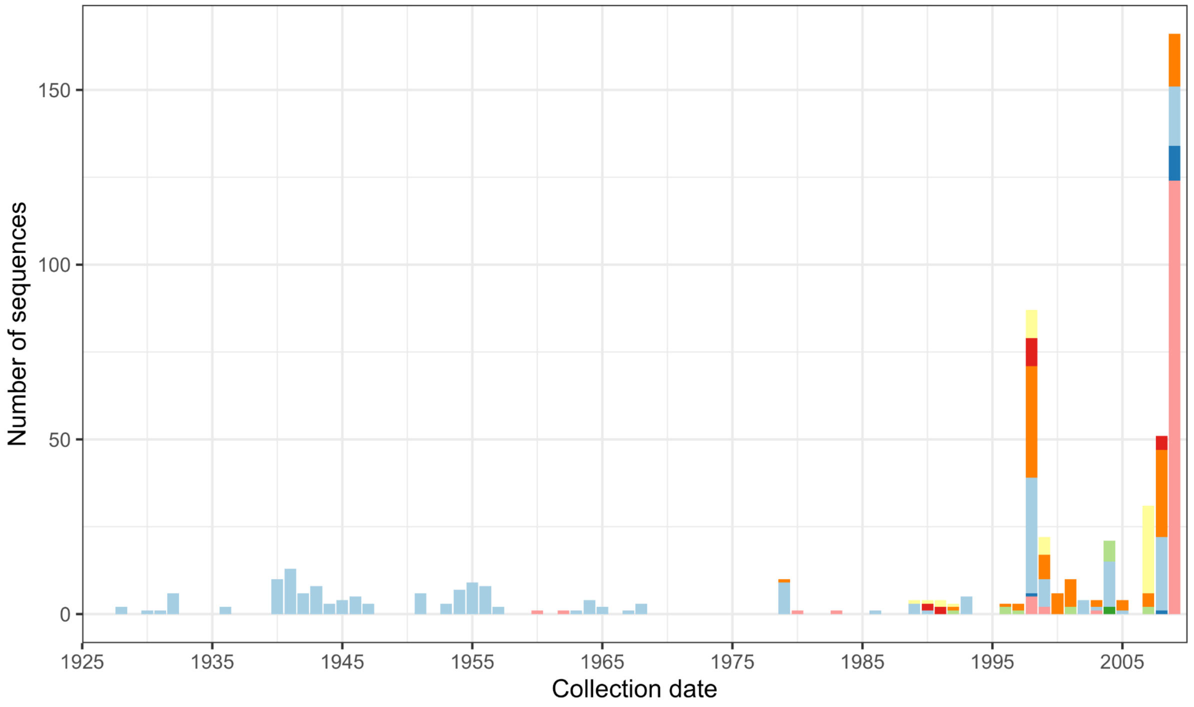


**Figure S1. Collection dates of the genomes. A)** The collection dates of the genomes stratified by year from 1995-2019 and colored by the location as indicated in the legend. **B)** Older isolates from 1925-2005.

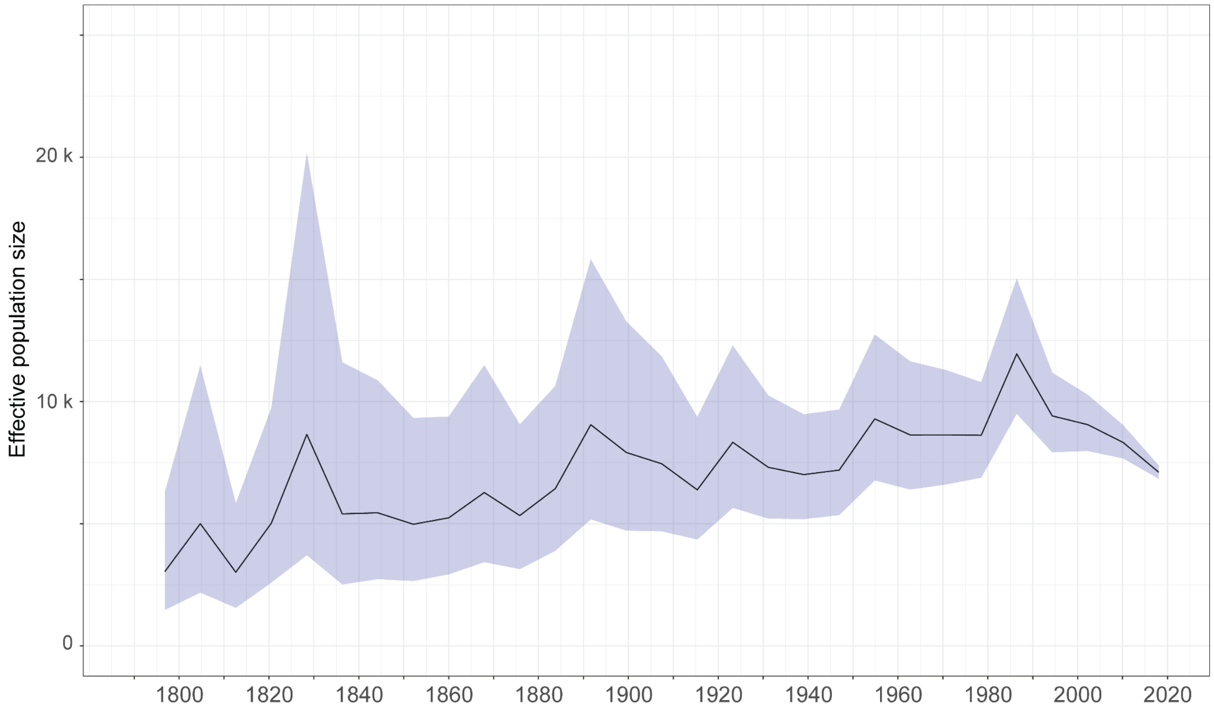


**Figure S2. Effective population size estimated from minimally biased collection of genomes.** As portions of the total sequence collection were not made up of randomly selected samples, the effective population size over time was estimated here from a dataset containing 4184 randomly sampled genomes. This dataset was made up of the genome collections from Norway, Victoria, the UK GoTG-trial dataset and the historic collection of gonococcal genomes from Denmark, the latter to provide temporal calibration points in the tree. A phylogeny was built using FastTree (Price, Dehal, and Arkin 2010) using the GTR substitution model. A dated tree was generated using BactDating (Didelot et al. 2018) and effective population size over time was reconstructed using skygrowth (Volz and Didelot 2018).

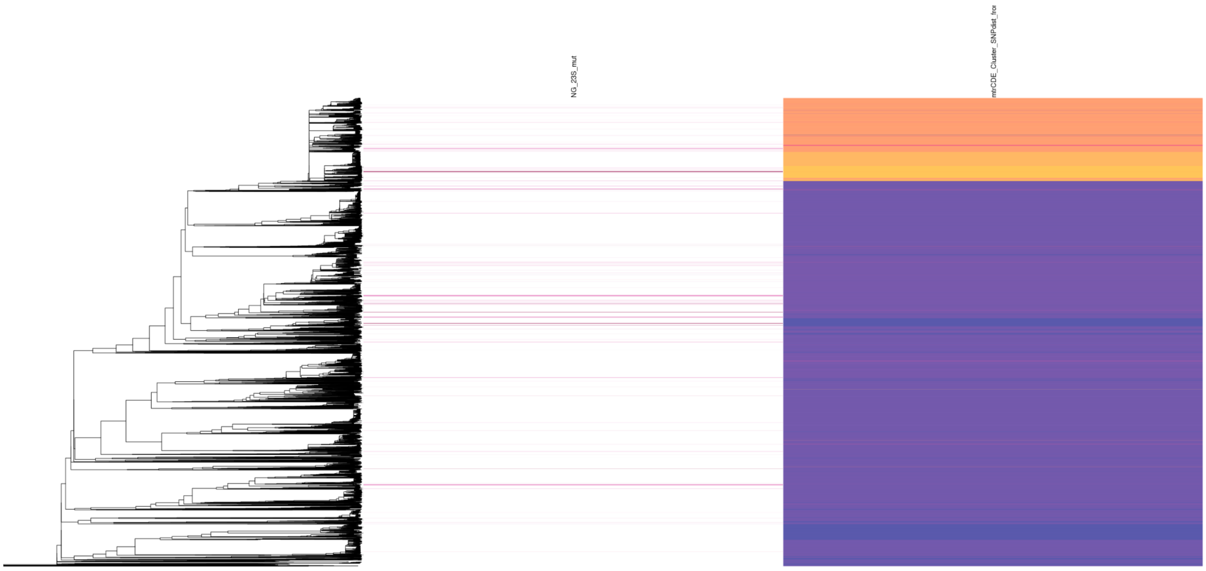

**Figure S3. *23S* and *mtr* alleles mapped to the dated phylogeny.** The annotation column to the left indicates the presence of 23S (*rrl*) mutations (A2059G, C2611T or both). The column on the right indicates the divergence of the *mtrCDE* operon relative to the allele in the FA1090 reference (yellow most diverged, blue least diverged).

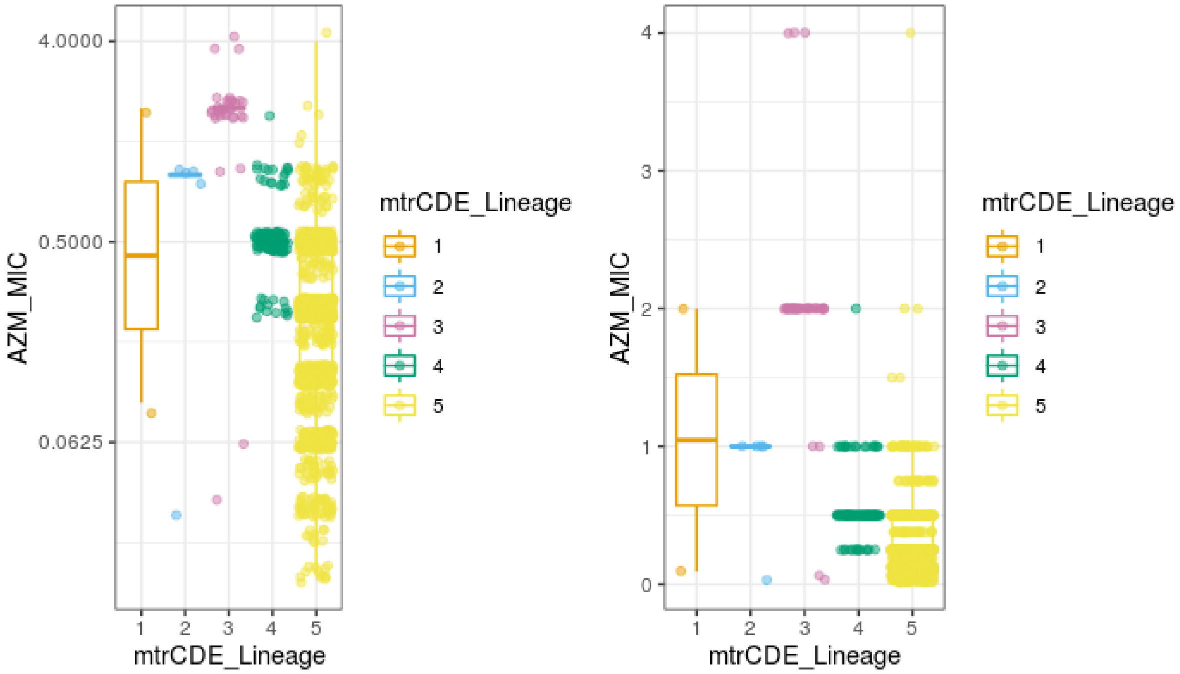


**Figure S4. AZM MICs stratified by *mtr* lineage as identified by FastGear.** Lineage 5 includes non-mosaic alleles, whereas lineages 1-4 represent different mosaic variants. Isolates harboring *23S* A2059G and/or C2611T mutations were excluded. *mtrCDE* lineage 5 corresponds to non-mosaic in Fig.2 in the main text.

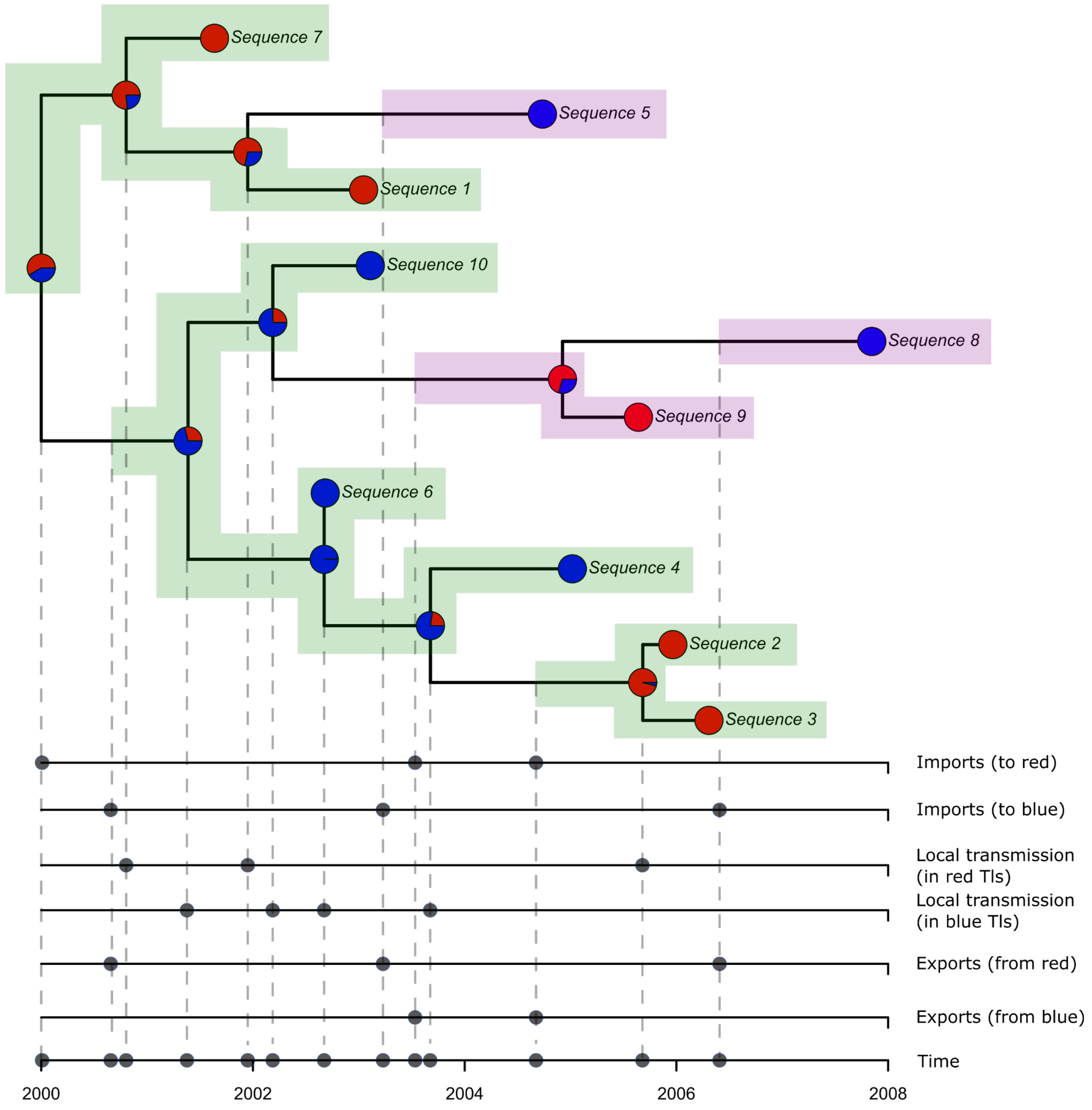


**Figure S5. How LineageHomology derives import, export and local transmission dates from geographically mapped phylogenies.** The green-shaded background represents transmission lineages and encompasses the tips that are included in the group. The probability of the locations is assumed to be obtained from an ancestral trait reconstruction and is shown by the fraction of the pie-charts filled with the relevant color. The transmission lineages are defined based on a 50 percent probability of the same location on every node within the TL. The pink-shaded background denotes singletons, which are unconnected to any other tips as per the rule that defines TLs. The areas showing TLs and singletons extend back to the importation date, which is the midpoint of the edge ancestral to the most recent common ancestor for the TLs and the midpoint on the ancestral edge leading to the first geographical transitions for singletons. Exports are set to the time at the midpoint of the branch that leads to a different geographical location from the TL. Branching events within TLs are used as estimates of local transmission. All types of events are projected down as points to individual axes to display the events with respect to the red or blue location in this example.

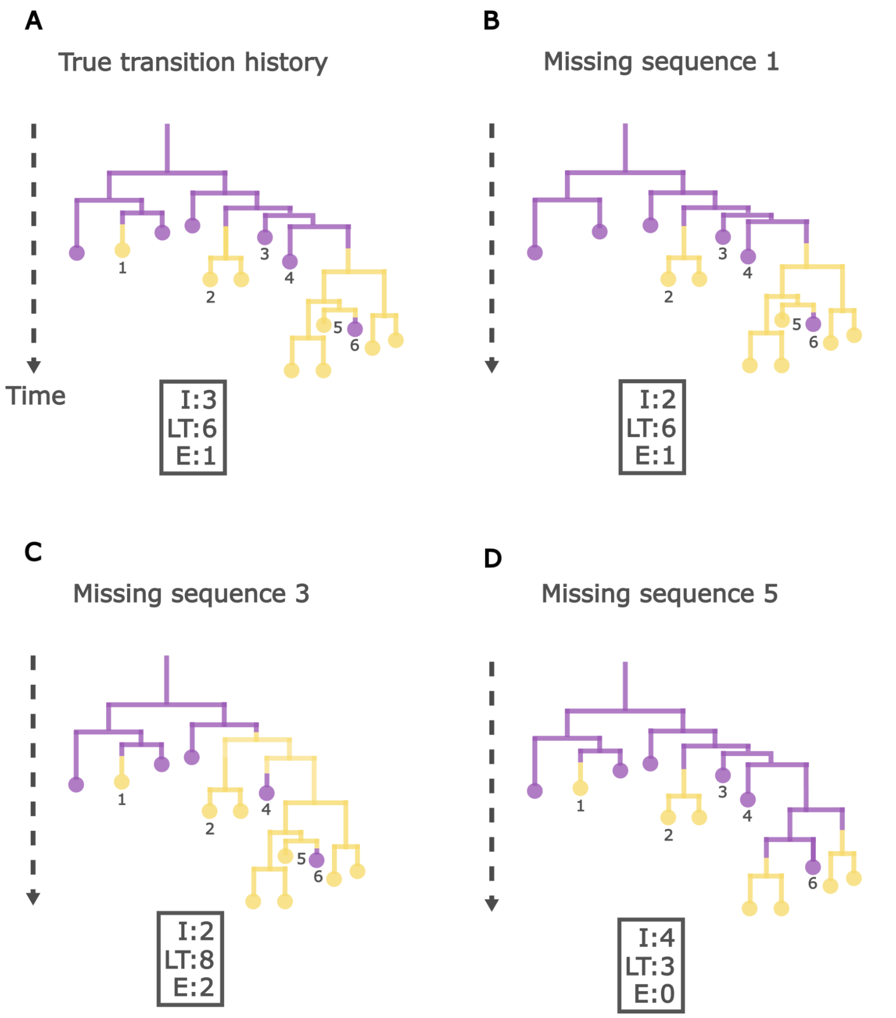


**Figure S6. The impact of collection bias on the estimates from LineageHomology with yellow as the location of interest. A)** The true geographical transmission history is shown with two locations (yellow and purple), where yellow represents the location of interest. Expected import (I), local transmission (LT), and export (E) events estimated by LineageHomology are indicated in the boxes. In this representation, there are three import events, two of which result in more than one local case in the yellow location. The procedure should correctly identify three importation events and six local transmission events (branching events within the transmission lineages). **B)** If surveillance fails to include sequence 1 in the genomic collection, it would result in one less estimated import event. **C)** Sequence 3 provides information about the true (purple) geographic state of the most recent common ancestor of both transmission lineages. If the collection does not include this sequence, both transmission lineages may collapse into one, resulting in an increase in local transmission events from six to eight, with one of these local transmissions estimated to result in an export event (sequence 4). **D)** Sequence 5 is important for the analysis of the geographical state of the ancestral branches of the transmission lineages. If we lose this sequence, the transmission lineage may split, leading to an increase in import events from three to four and a decrease in local transmissions from six to three. **In general,** we expect larger transmission lineages to have a higher detection probability (since the detection of one infected in a location will likely increase the detection of others infected by the same strain). Hence we expect an increased probability of detection of larger transmission lineages and a lower probability of detection of singletons. Therefore, we expect scenarios **B** and **C** to be common in collections with high sampling densities in the location of interest, and **D** to be less common. This results in a bias towards fewer import events and more local transmission events when using the LineageHomology approach. The figure is inspired by the arguments in the supplementary text of (du Plessis et al. 2021).

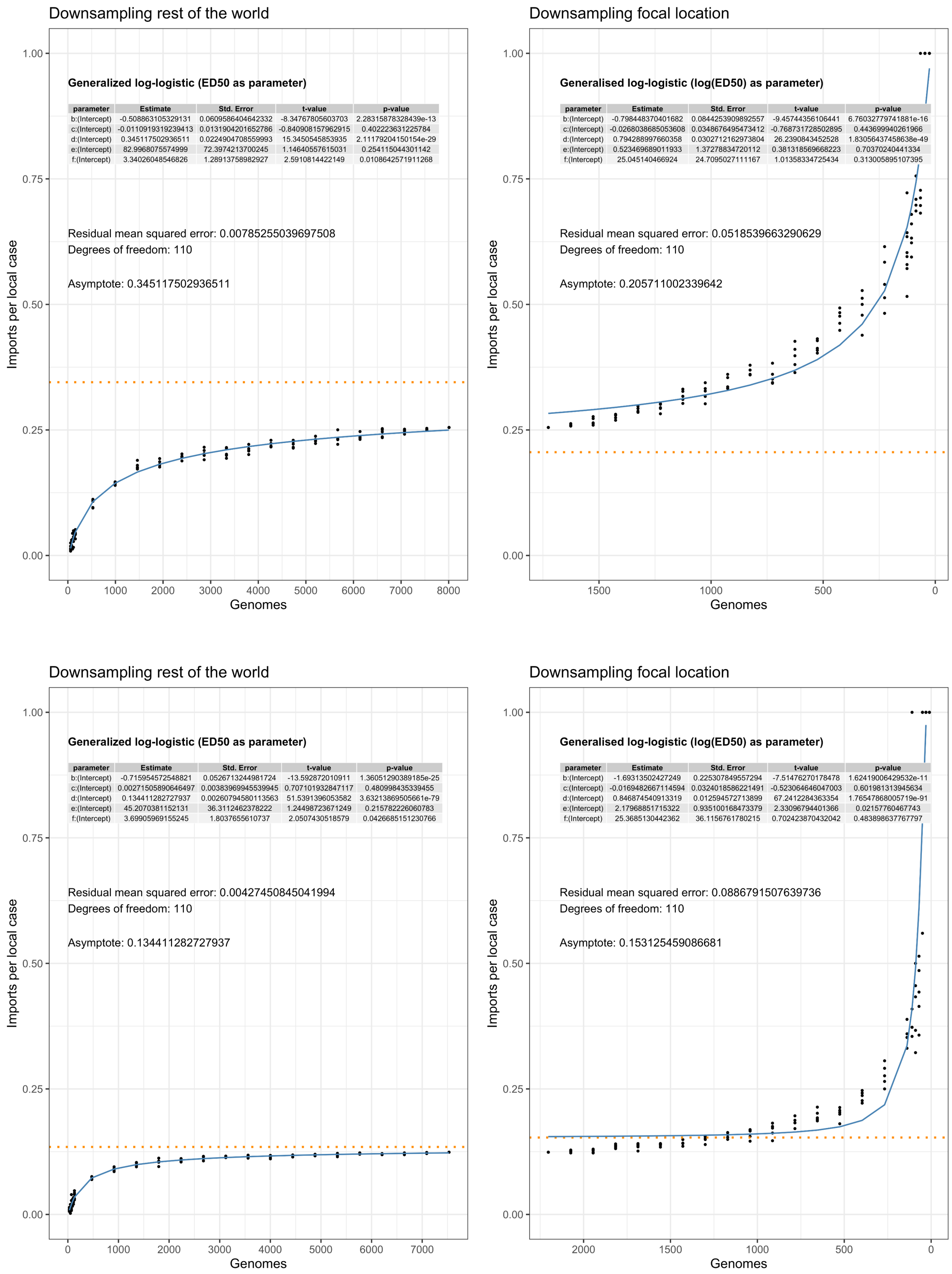


**Figure S7. Import fraction asymptote for Norway (top) and Victoria, Australia (bottom).** The panels on the left side display the results from upsampling (effectively downsampling tips of the phylogeny) the contextual samples from outside the focal location (Rest of World, ROW). The panels on the right illustrate the estimates based on upsampling within the focal location. The inset text specifies the name of the best-fitting asymptotic function, with the estimated parameters tabulated. The orange line represents the estimated asymptote, while the blue line depicts the curve predicted by the model. Additional details are provided in the main text.

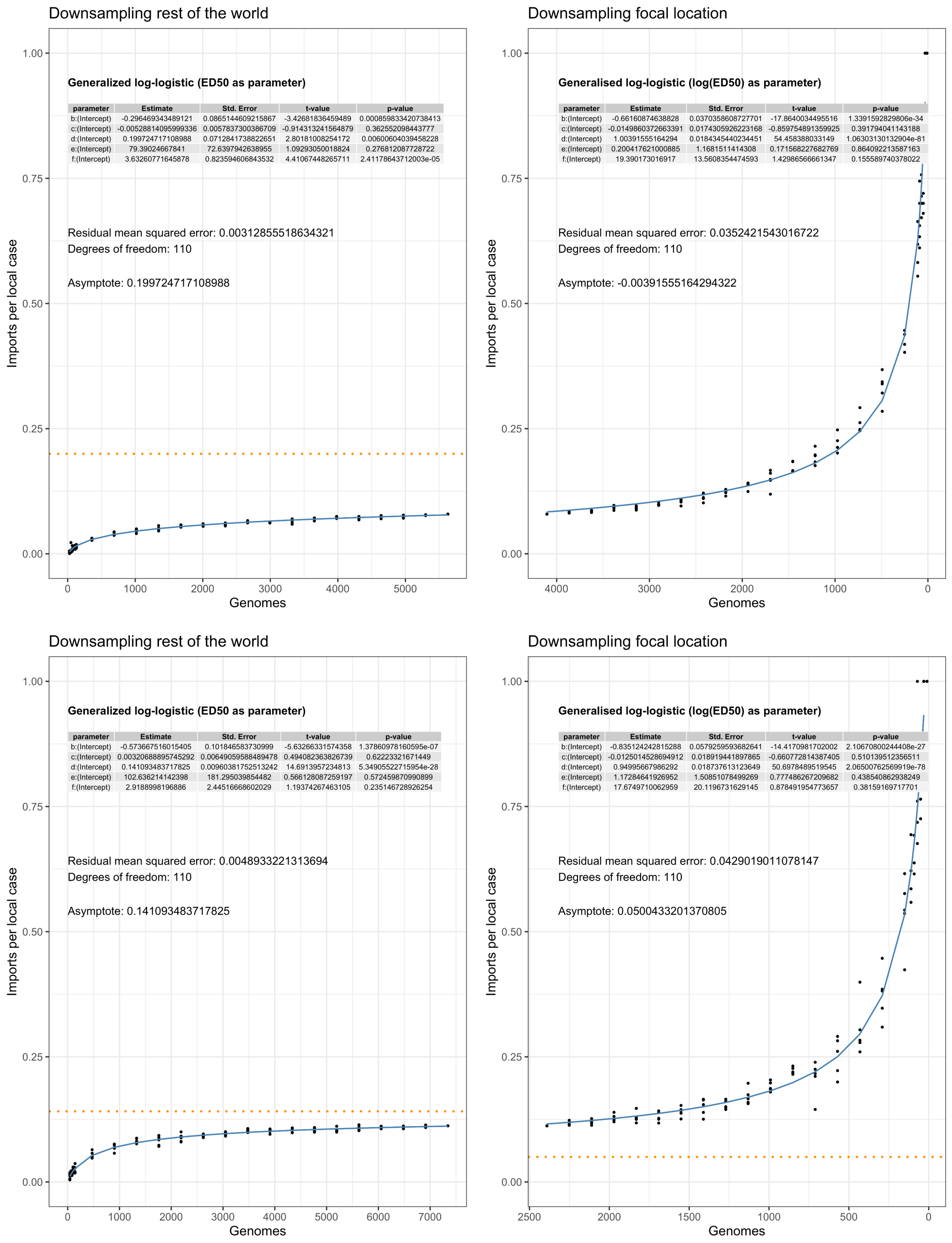


**Figure S8. Import fraction asymptote for Europe (top) and the USA (bottom).** See Figure S7 and the main text for details.

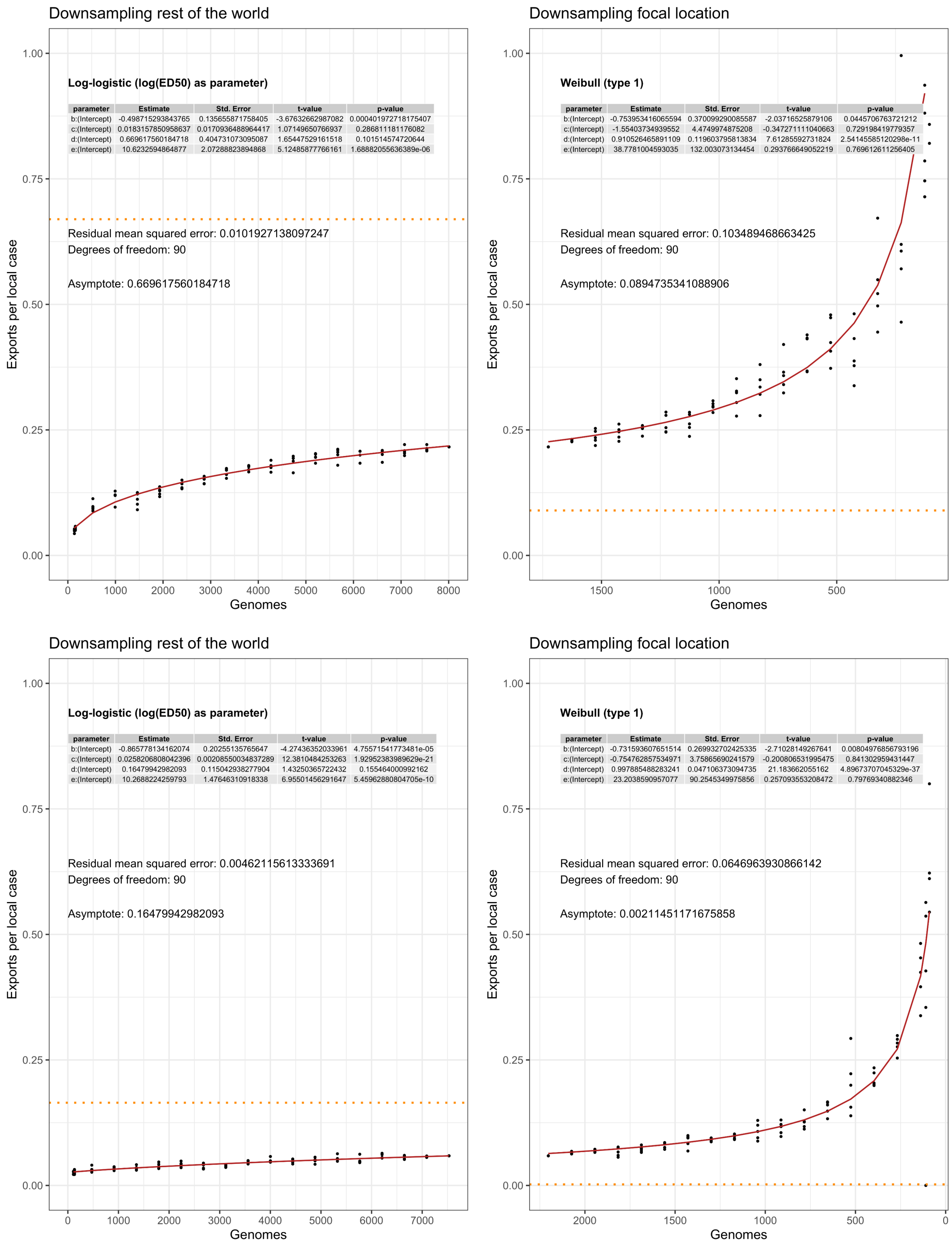


**Figure S9. Export fraction asymptote for Norway (top) and Victoria, Australia (bottom).** See Figure S7 and the main text for details.

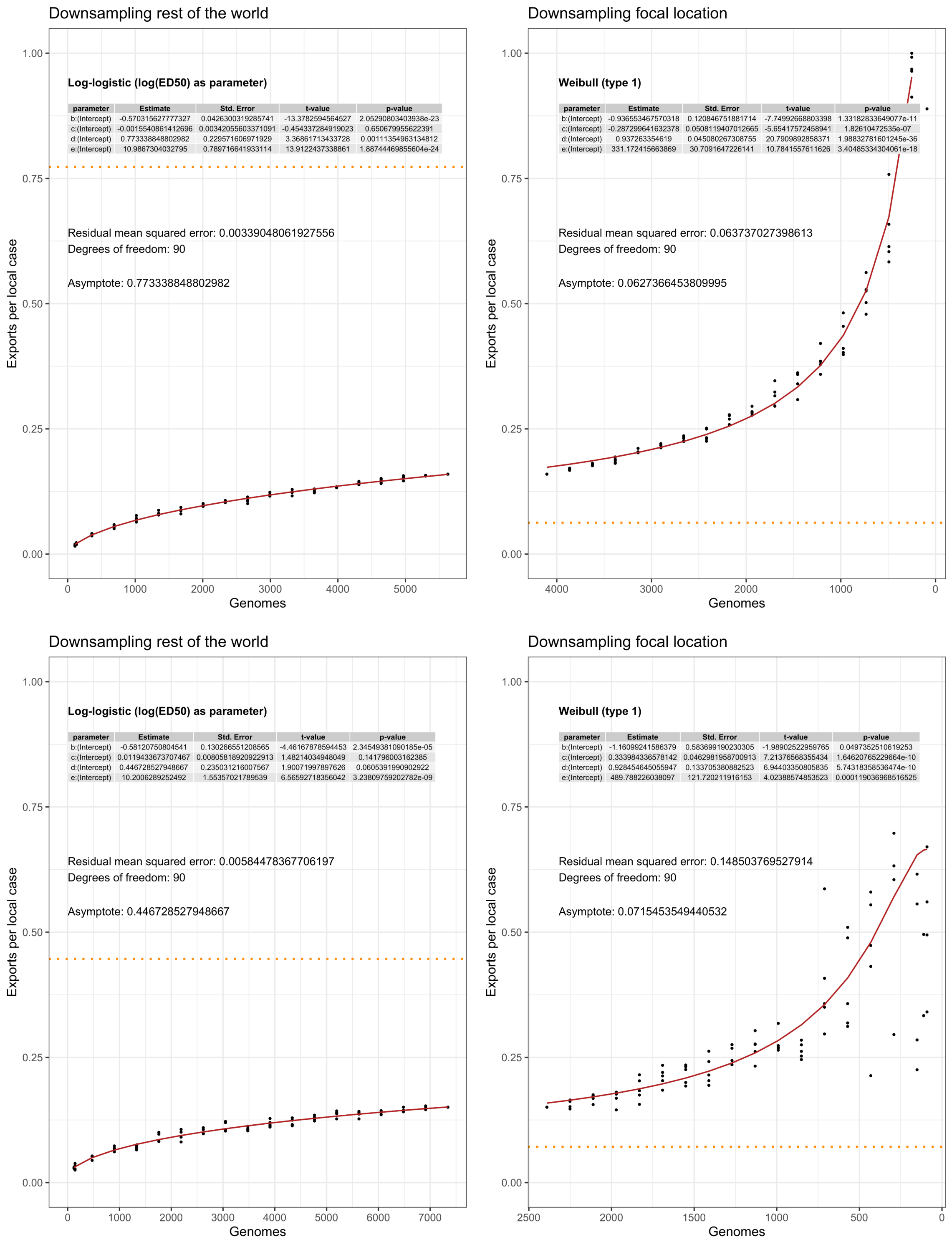


**Figure S10. Export fraction asymptote for Europe (top) and the USA (bottom)**. See Figure S7, and the main text for details.

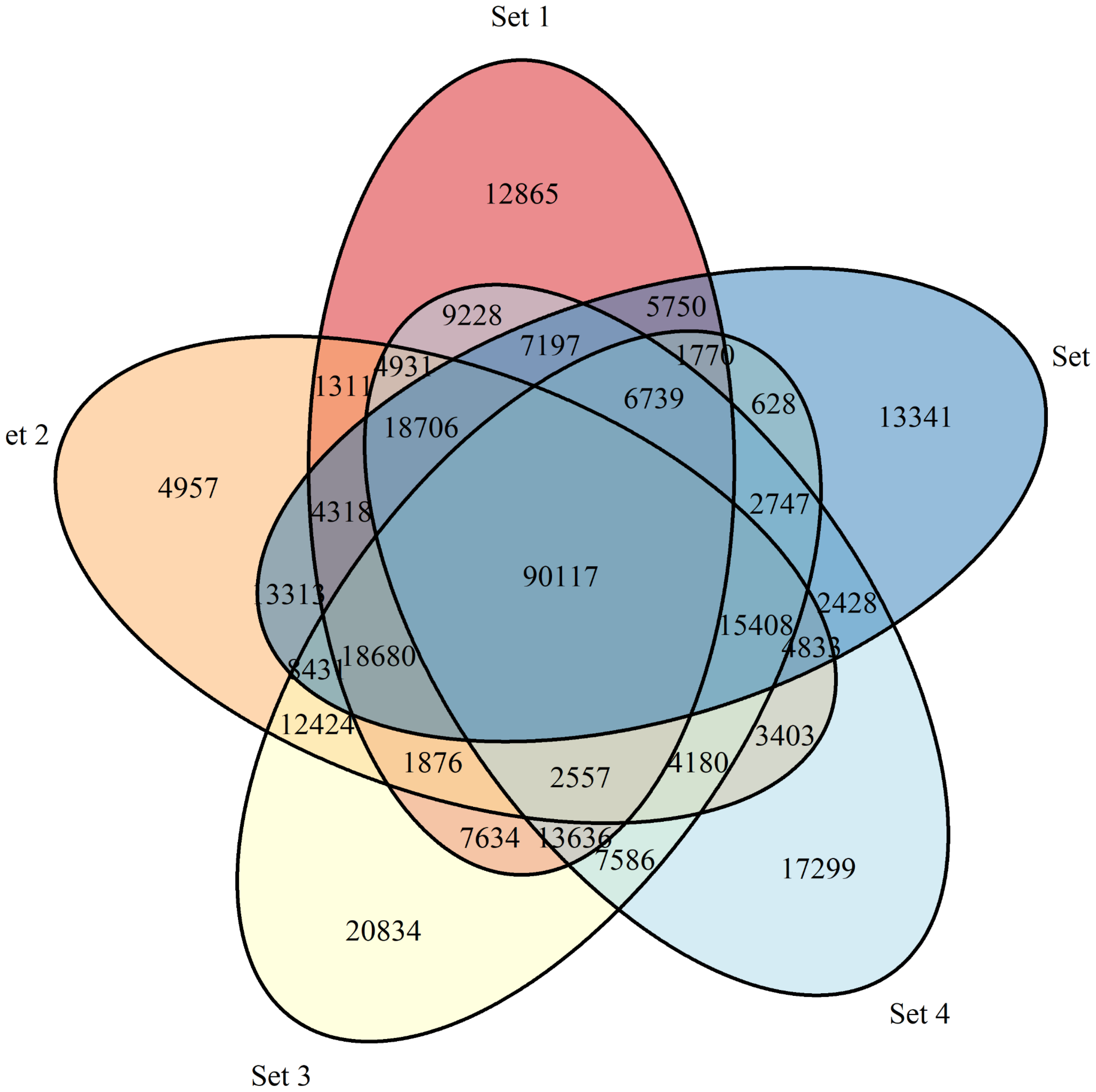


**Figure S11. Details on recombination filtering.** Venn diagram illustrating the overlap between identified highly recombined regions across five genome subsets (see main text for details).

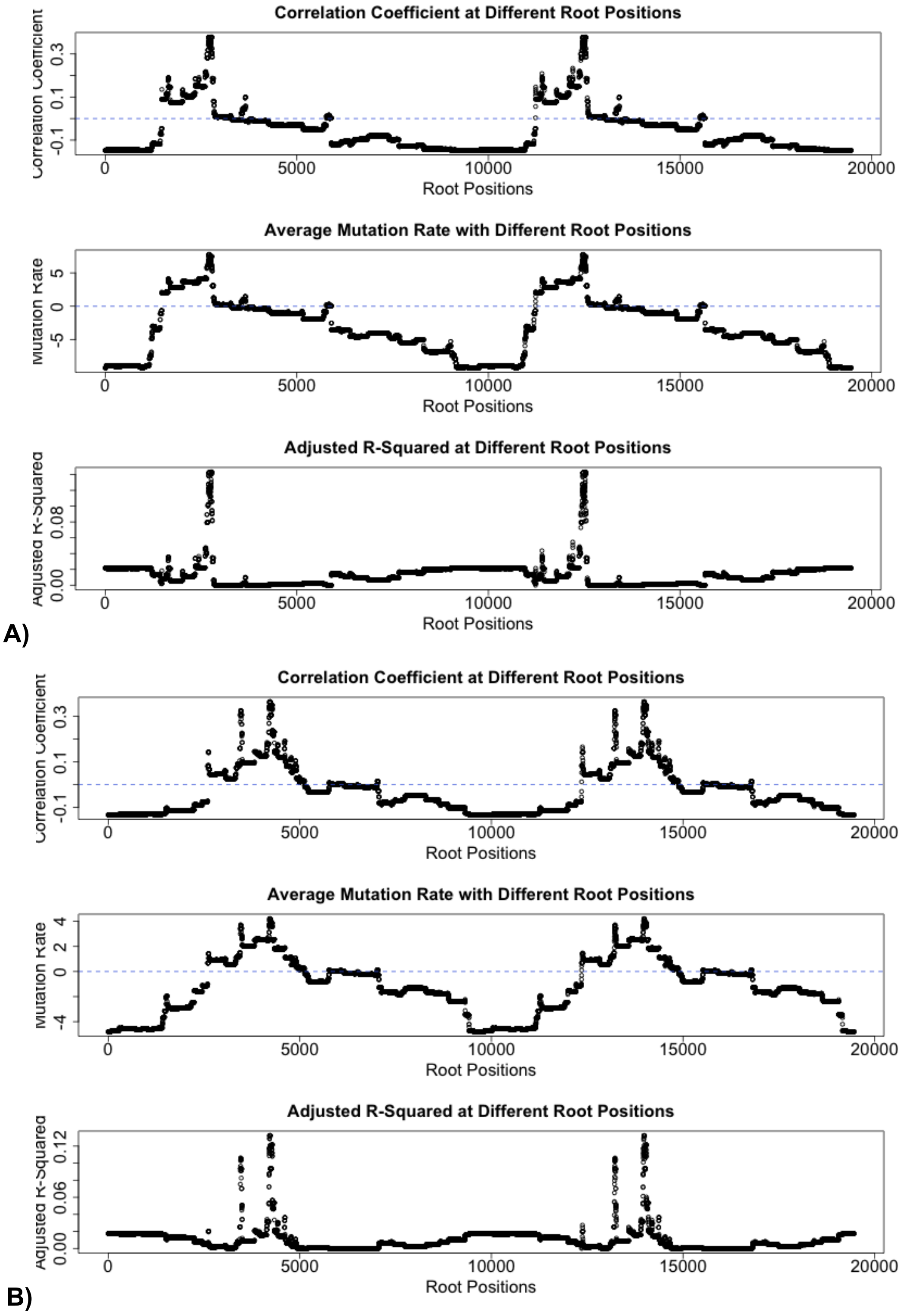


**Figure S12. Temporal signal analysis for different root positions.** (A) Root positions in the phylogeny without masked recombination. (B) Root positions in the recombination-masked phylogeny. Top panel: Correlation of root-to-tip distance with collection dates. Middle panel: Estimated average mutation rate per genome per year at each root position. Bottom panel: Adjusted R-squared values for each root position's temporal signal. The pattern is duplicated in all figures because the analysis first considers leaves as possible root positions (position1 to 9732) and then internal nodes (position 9732-19463).

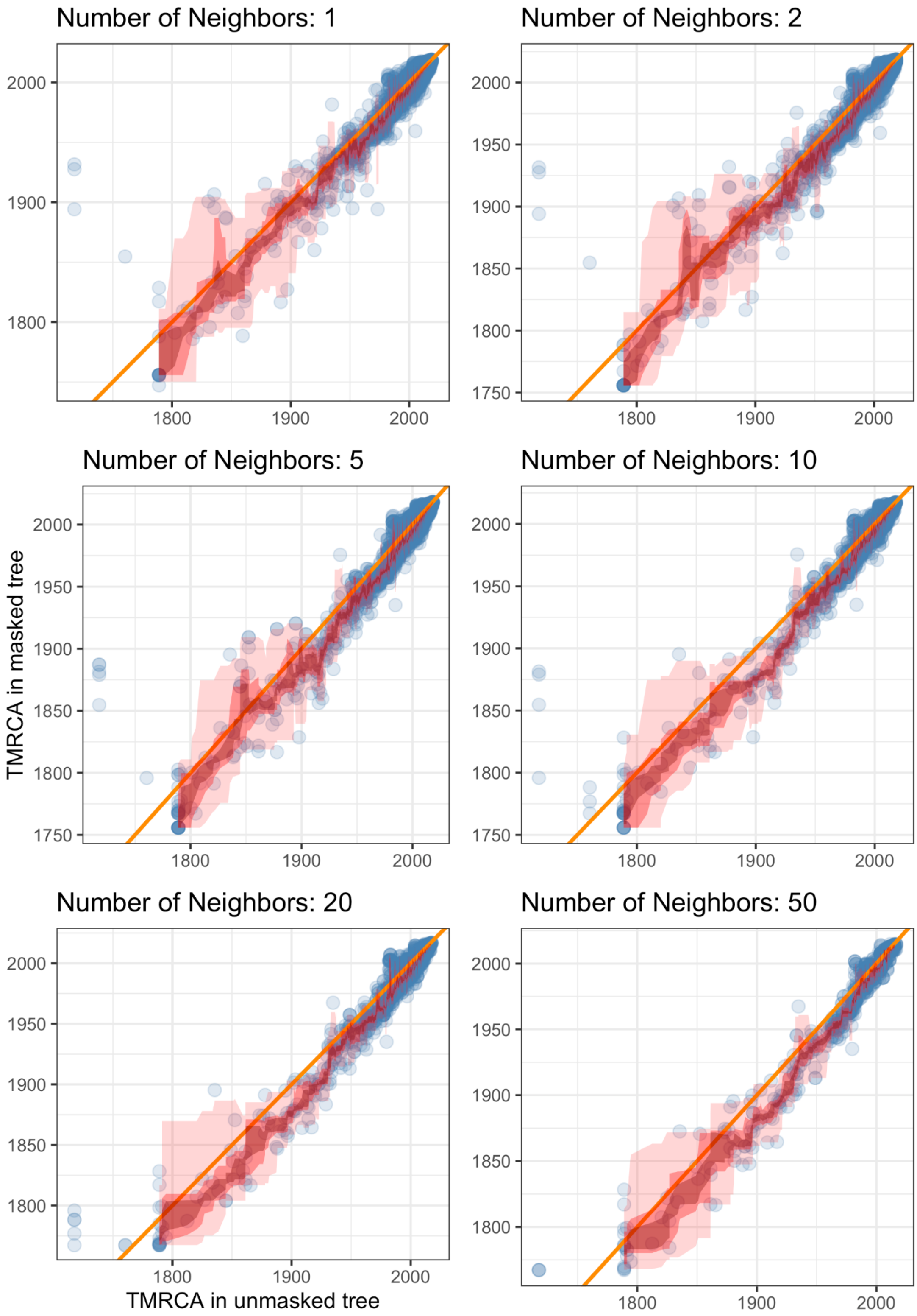


**Figure S13. The estimated time to the most recent common ancestor (TMRCA) for recombination-masked and unmasked trees.** This figure illustrates the differences in TMRCAs of leaves and their 1, 2, 5, 10, 20, and 50 closest neighbors in the recombination-masked and unmasked trees. The analysis indicates that the recombination-masked trees generally estimated older TMRCAs for most points. Further details of the methods for this analysis are available in the Supplementary Methods section.

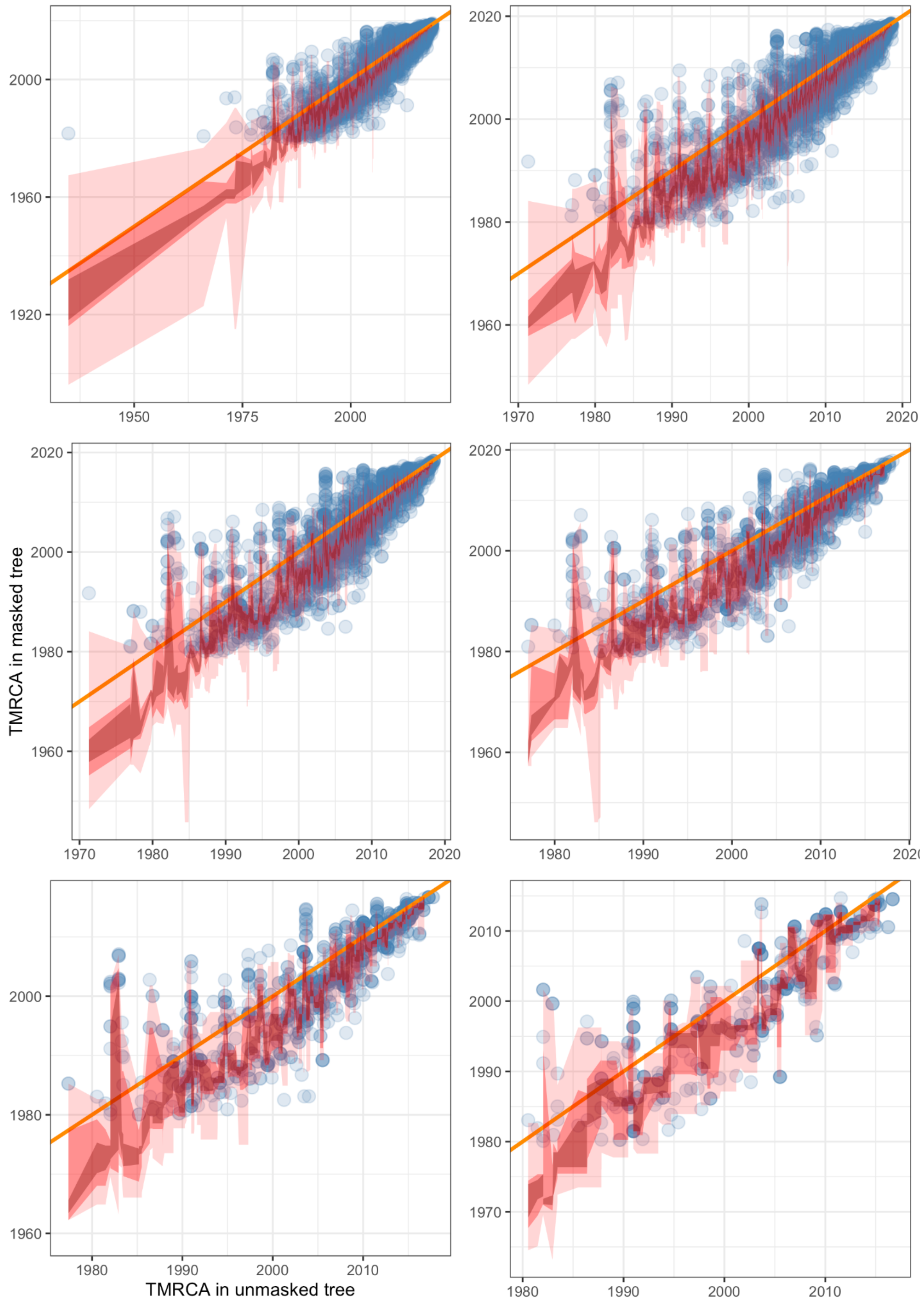


**Figure S14. The estimated time to the most recent common ancestor (TMRCA) for recombination-masked and unmasked trees between 1980 to 2018.** This figure shows the evolution of differences between TMRCAs in recombination-masked and unmasked trees over time, showing a trend of diminishing differences between TMRCAs as we approach the present. The methodology used for this analysis can be found in the Supplementary Methods section.

#### References


Didelot, Xavier, Nicholas J. Croucher, Stephen D. Bentley, Simon R. Harris, and Daniel J. Wilson. 2018. “Bayesian Inference of Ancestral Dates on Bacterial Phylogenetic Trees.” *Nucleic Acids Research* 46 (22): e134.

Plessis, Louis du, John T. McCrone, Alexander E. Zarebski, Verity Hill, Christopher Ruis, Bernardo Gutierrez, Jayna Raghwani, et al. 2021. “Establishment and Lineage Dynamics of the SARS-CoV-2 Epidemic in the UK.” *Science* 371 (6530): 708–12.

Price, Morgan N., Paramvir S. Dehal, and Adam P. Arkin. 2010. “FastTree 2--Approximately Maximum-Likelihood Trees for Large Alignments.” *PloS One* 5 (3): e9490.

Volz, Erik M., and Xavier Didelot. 2018. “Modeling the Growth and Decline of Pathogen Effective Population Size Provides Insight into Epidemic Dynamics and Drivers of Antimicrobial Resistance.” *Systematic Biology* 67 (4): 719–28.
